# Microbiome plasticity, not gut morphology, is linked to amphibian larval performance under elevated temperatures and low food quality

**DOI:** 10.64898/2026.07.25.740725

**Authors:** Katharina Ruthsatz, Myra C. Hughey, Maline Türk, Marjoriane de Amaral, Julian Glos, Paula C. Eterovick

## Abstract

In many ecosystems, anthropogenic warming is reshaping thermal regimes, leading to resource quality declines and imposing a dual constraint for ectotherms: elevated metabolic demand coupled with reduced assimilable energy. We tested whether plasticity in gut morphology and gut microbiome can buffer amphibian larvae against these concurrent stressors. Common frog (*Rana temporaria*) tadpoles were reared at two temperatures (18 vs. 24.5°C) crossed with three food-quality treatments (low, medium, high). We quantified growth and developmental rates, critical thermal limits (CT_max_, CT_min_), gut morphology (mass, relative length), and gut bacterial diversity and composition, together with predicted functional pathways. Warming accelerated growth and development and increased CT_max_. Food quality increased growth and development, with temperature-dependent effects on developmental rate and CT_max_. Gut mass declined at higher temperature and low-quality diets, but relative gut length showed only modest diet effects and no temperature dependence. Bacterial community composition and structure shifted with temperature and food quality. Predicted pathways suggest functional reconfiguration under warming and low food quality, consistent with sustaining energy acquisition and mitigating metabolic and oxidative stress. Together, these results implicate microbiome plasticity, rather than gut morphological plasticity, as a candidate mechanism supporting larval performance and heat-tolerance acclimation under warming and low food quality.

## 1. Introduction

Global change is reshaping environments through shifts in temperature regimes and alterations in resource availability. Rising mean temperatures, greater thermal variability, and more frequent extremes such as heat waves or cold spells (IPCC 2023) interact with declining food quality in many ecosystems (Paerl & Huisman 2009; Yan et al. 2024), creating major challenges for organisms to acquire sufficient energy to sustain growth, performance, and survival. Ectothermic species are particularly vulnerable to these changes, as their body temperature and thus all physiological processes depend directly on environmental temperature (Hochachka & Somero 2002). Among the physiological rates shaped by ambient temperature, metabolic rate is particularly important, as it sets the organism’s energetic demands and is highly sensitive to thermal change (Norin & Metcalfe 2019). At warmer temperatures, ectotherms must therefore meet elevated energetic requirements through greater food intake and efficient assimilation. Yet, digestive performance is itself temperature dependent (Fontaine et al. 2018), constraining the capacity of organisms to match elevated demands with effective energy acquisition. If this balance fails, energetic shortfalls can cascade across levels of biological organization, reducing growth and development, impairing performance, altering behavior, and ultimately lowering survival. As multiple environmental stressors can interact to exacerbate these constraints (Sinai et al. 2024), identifying mechanisms that allow ectotherms to adjust and maintain function is fundamental for predicting their resilience to global change.

Digestive performance is not fixed and can be adjusted through multiple physiological pathways. Gut microbial communities, for example, often shift in composition in response to environmental stressors such as altered diet, pollutants/toxins, or temperature change (diet: Youngblut et al. 2019; Hardison & Eliason 2024; temperature: Sepulveda & Moeller 2020; Huus & Ley 2021; toxins: Jin et al. 2017; Eterovick et al. 2024), enabling hosts to improve digestive efficiency or exploit alternative nutrient sources through detoxification or fermentation (Mackie 2002; McFall-Ngai et al. 2013; Kohl et al. 2014). Similarly, plasticity in gut morphology represents a widespread adaptive response across vertebrate taxa (Secor 2011). Gut elongation, for example, can increase retention time and facilitate microbial processing of otherwise indigestible material (Yang & Joern 1994). These adjustments of host tissues and microbial symbionts may interact, with shifts in microbial function complementing structural modifications of the gut to optimize digestion under environmental stress. By improving nutrient assimilation under constrained food availability, these mechanisms may buffer hosts against energetic shortfalls and support developmental plasticity, behavioral flexibility, and enhanced thermal tolerance. Yet, despite abundant evidence for stressor-induced changes in gut microbiota and morphology separately, little is known about how these responses interact to sustain host physiology under multiple co-occurring stressors.

Another key trait determining how organisms cope with temperature fluctuations is thermal tolerance, which defines the limits of physiological function and survival (Angilletta et al. 2002). Performance generally follows a thermal performance curve, declining steeply near the critical thermal minimum (CT_min_) and maximum (CT_max_; Sinclair et al. 2016). Both limits are relevant under climate change: warming and more frequent heat waves increase demands on heat tolerance, while late frosts or false springs may expose temperate ectotherms to sudden cold spells (Ault et al. 2013; Gunderson & Stillman 2015). Importantly, these traits are not fixed but vary with several extrinsic and intrinsic factors such as phylogeny, ontogeny, acclimation/thermal history, and nutritional state (Ruthsatz et al. 2022a, 2024).

Mounting evidence suggests that gut microbial communities can modulate host thermal tolerance (Fontaine & Kohl 2023; Hardison & Eliason 2024). Experimental studies across invertebrates and vertebrates show that temperature-induced shifts in microbiota composition can alter heat tolerance (Sepulveda & Moeller 2020), although the underlying mechanisms remain poorly resolved, particularly in vertebrates (Fontaine & Kohl 2023). Correlative evidence suggests that such shifts may improve nutrient assimilation, increase energy yield from complex diets, or alter host metabolism through microbial metabolites such as short-chain fatty acids (Warne & Dallas 2022; Dallas et al. 2024). These metabolites may enhance antioxidant capacity and buffer heat-induced oxidative stress, thereby supporting heat tolerance (Fontaine et al. 2022; Sokolova 2023). The gut microbiome may therefore support thermal performance indirectly by helping maintain host energy balance under elevated metabolic costs (Fontaine et al. 2018, 2022; Dallas et al. 2024). At the same time, changes in digestive morphology including gut elongation under low-quality diets, may further interact with microbial communities to influence thermal performance.

Despite their potential importance as coping mechanisms, plasticity in digestive morphology and gut microbiota have rarely been studied in conjunction with thermal physiology, and to our knowledge no study has yet integrated food quality, gut morphology, and microbial composition with measures of thermal tolerance under shifting thermal regimes. This leaves a critical gap in our understanding of how interacting environmental drivers shape multiple axes of physiological plasticity and, ultimately, organismal resilience to global change. Addressing this gap raises a key question: *do organisms have the “guts” to cope with climate change?* – that is, *can plasticity in gut morphology and microbial symbionts help maintain energy acquisition under simultaneous shifts in temperature and food quality, thereby supporting sufficient thermal tolerance*?

Amphibians provide a powerful system for addressing this mechanistic question. Their aquatic larvae have limited dispersal and are highly sensitive to changes in temperature and resource quality (Álvarez & Nicieza 2002; Kohl & Yahn 2016). Amphibian larvae also exhibit substantial plasticity in growth, developmental timing, gut morphology, microbiome composition, and stress physiology (Relyea & Auld 2004; Gomez-Mestre et al. 2010; Fontaine & Kohl 2020), making them well suited for testing how digestive and thermal responses interact to sustain performance under environmental change. This is particularly relevant because amphibians are the most threatened vertebrate group globally, with climate change compounding declines driven by habitat loss, disease, and pollution (Luedtke et al. 2023). Because microbiomes can respond rapidly to stressors and alter host metabolic functions, they may increase amphibian adaptive capacity under such conditions (Louca et al. 2018; Voolstra & Ziegler 2020; Fontaine & Kohl 2023).

Here, we used European common frog (*Rana temporaria*) larvae to test whether plasticity in gut morphology and the microbiome facilitates responses to concurrent variation in food quality and temperature. In a fully factorial design, larvae were reared at three food-quality levels and two temperatures. We quantified thermal tolerance (CT_min_ and CT_max_), gut morphology (length and mass), microbial composition, and predicted functional pathways. We also tested whether diet- and temperature-induced shifts in gut morphology, indicator taxa, and metabolic potential covaried with growth and thermal tolerance. By integrating these potentially complementary dimensions of phenotypic plasticity, our study evaluates how amphibian larvae cope with simultaneous shifts in food quality and temperature, offering new insights into the mechanisms that shape ectotherm resilience under climate change.

## 2. Materials and Methods

### 2.1 Study species and field sampling

We used the European common frog (*Rana temporaria*) as a model species because of its high plasticity in metamorphic and physiological traits in response to environmental variation, including temperature and food quality (Ruthsatz et al. 2019; Sinai et al. 2024). Five egg clutches were collected on 25 March 2023 from a breeding pond near Braunschweig, Lower Saxony, Germany (52.328° N, 10.582° E), and transported to Technische Universität Braunschweig. Clutches were maintained separately until hatching, after which larvae were reared at 18 °C under a 14:10 h light cycle until Gosner stage 25.

### 2.2 Experimental design and animal husbandry

From each clutch, 84 larvae were haphazardly selected and assigned to a fully crossed design comprising two rearing temperatures (18 and 24.5 °C) and three food-quality treatments (low, medium, and high; Section 2.3). This resulted in 70 individuals per treatment combination and a total sample size of 420 larvae. Larvae were housed individually in 1.2-L containers filled with 1 L of dechlorinated tap water. The 18 °C treatment was maintained in a climate chamber, whereas the 24.5 °C treatment was established using heated water baths located within the same chamber. Water temperature was maintained at 18 ± 0.1 or 24.5 ± 0.4 °C, respectively. Water was replaced every third day. After five weeks of exposure, subsets of larvae were allocated to the different measurement endpoints described in Section 2.4. Further details of the collection, acclimation, temperature-control system, and husbandry procedures are provided in Electronic Supplementary Material.

### 2.3 Experimental feeding regime

Larvae were reared under low-, medium-, or high-quality diets differing in ingredient diversity, protein content, and energy density. The low-quality diet consisted of organic barley grass powder (*Hordeum vulgare*; NaturaleBio, Germany: 3% lipid, 11% carbohydrate, and 32% protein), containing leaves from a single plant species and comparatively little protein. The high-quality diet consisted of powdered fish food (Sera micron, Sera, Germany: 7.2% lipid, 10.3% carbohydrate, and 56.6% protein), comprising algae, zooplankton, and plant and animal products and providing more protein of both vegetal and animal origin. The medium-quality diet was a 50:50 mixture of both food types. See Electronic Supplementary Material for information on dietary energy content and stable isotope analysis.

### 2.4 Allocation to measurement sets

The study integrated three complementary measurement sets within a single experimental design: (i) thermal tolerance, (ii) gut morphology, and (iii) gut microbiome composition. When larvae reached pre- to early pro-metamorphic stages (Gosner 27–36), individuals were sampled for different endpoints (N=388). Specifically, 105 larvae were used for thermal tolerance assays, 170 for gut morphology, and 53 for microbiome analyses. Sixty additional larvae were diverted to a heatwave simulation experiment reported elsewhere (Eterovick et al. 2026).

### 2.5 Sampling and metamorphic measurements

At the conclusion of the experiment, all surviving larvae (N = 388) were euthanized with 2 g L⁻¹ tricaine methanesulfonate (MS-222; Sigma-Aldrich). Thirty-two larvae died over the course of the experiment. For each individual, total length (TL), snout–vent length (SVL), body mass, and developmental stage (*sensu* Gosner 1960) were recorded. Body mass had also been measured at Gosner stage 25. SVL was measured with a caliper to the nearest 0.5 mm, and body mass with a precision balance (0.001 g; Sartorius A200 S, Germany) after blotting dry. Growth rate (mg d⁻¹) was calculated as the difference in body mass between stage 25 and final sampling, divided by days elapsed.

#### 2.5.1 Thermal tolerance measurements

Critical thermal minimum (CT_min_) and maximum (CT_max_) were measured in subsets of larvae as proxies for thermal tolerance (N = 49 and 50, respectively, as 6 larvae died during the measurements). We followed the dynamic method (Lutterschmidt & Hutchison 1997), with the endpoint defined as loss of righting response (Wu & Kam 2005). To assess this, the larvae were gently flipped onto their backs in water using a probe (Sinai et al. 2024).

Each larva was placed individually in a 250-mL beaker containing 200 mL of aerated water positioned in a temperature-controlled water bath. Initial water temperature matched the larva’s rearing temperature (18 or 24.5 °C). For CT_min_ trials, water temperature was decreased at 0.5 °C min⁻¹ until the endpoint; for CT_max_ trials, water temperature was increased at 0.1 °C min⁻¹. After reaching CT_min_ or CT_max_, larvae were gently transferred to a beaker containing water at a temperature halfway between their acclimation temperature and the measured critical temperature. The beaker was placed in a water bath set to the appropriate rearing temperature to allow gradual equilibration.

#### 2.5.2 Intestinal measures

For gut morphology analyses, guts were dissected from each larva, while tail and body remains were placed in sterile 1.5-mL tubes, snap-frozen in liquid nitrogen, and stored at −20 °C. Guts were fixed in 1 mL of 4% paraformaldehyde for ≥48 h to preserve structural integrity, then uncoiled, photographed with a digital microscope, and measured in ImageJ. Gut length was measured from the posterior end of the *manicotto glandulare* to the vent, and gut mass, including contents, was recorded after blotting dry using a precision balance (0.001 g; Sartorius A200 S, Germany).

#### 2.5.3 Microbiome analyses

To assess bacterial community composition, guts were removed immediately after euthanasia and stored at −80 °C until processing. DNA was extracted using the QIAmp Fast DNA Stool Mini Kit (QIAGEN), and the V4 region of the 16S rRNA gene was amplified with primers 515F (5′-GTGCCAGCMGCCGCGGTAA-3′) and 806R (5′- GGACTACHVGGGTWTCTAAT-3′; Caporaso et al. 2011), using unique forward and reverse tag combinations for each sample. Negative controls were included for each extraction day (n = 5) and PCR run (n = 2). Known microbial community and DNA standards (Zymo Research Europe) served as positive controls for DNA extraction and PCR precision, respectively. PCR products were pooled, separated on a 2% agarose gel, and the target 251-bp fragment was purified using the Monarch DNA Gel Extraction Kit (New England BioLabs GmbH, Germany). Purified DNA was quantified with Qubit™ (Invitrogen) and sequenced at the Leibniz Institute DSMZ using an Illumina MiSeq 500 platform (paired-end 2 × 250 bp, v2 chemistry).

### 2.6 Bioinformatic analyses, alpha and beta diversity estimations

Sequences were imported into QIIME2 (Bolyen et al. 2019), denoised, quality filtered (median Illumina Quality Score above 30), and trimmed to 250 bp. From the initial 1,298,311 paired sequences, 74,623 were retained, with sequencing depth per sample ranging from 257 to 5,441 reads. All negative controls were excluded during quality filtering.

The Greengenes (16S rRNA) reference database (version gg-13-8; McDonald et al. 2012) was used to generate a phylogenetic tree in QIIME2 following Estaki et al. (2020). To classify the filtered sequences, a classifier was trained with the bespoke method using reference sequences, taxonomy and sequence weights from animal proximal gut (Silva release 138.1, 515F/806R) available at *github.com/BenKaehler/readytowear* (Kaehler et al. 2019). Reads obtained from the positive controls were individually checked for taxonomy with BLAST (National Center for Biotechnology Information; Sayers et al. 2025) instead of submitted to the classifier, as they are not samples from animal proximal gut.

We identified 157 unique amplicon sequence variants (ASVs; 13–56 per sample). No ASV represented <0.005% of reads retained after quality filtering, making the inclusion of amplification errors unlikely (Bokulich et al. 2013). Because many reads were lost during pairing, probably owing to low sequence quality, we estimated sample coverage and assessed curve stabilization by rarefaction using 150 evenly spaced x-axis points and 200 simulations in iNEXT (Chao et al. 2014). Coverage approached one and diversity curves stabilized for all samples, indicating adequate representation of microbiome diversity despite low sequencing depth. Bacterial alpha diversity was calculated as Hill numbers of order q = 1, based on the Shannon diversity index, using the R package iNEXT (Chao et al. 2014; Alberdi & Gilbert 2019). Hill numbers are robust to variation in sample depth and represent the number of equally abundant taxonomic entities producing the same diversity value (Hill 1973; Alberdi & Gilbert 2019). Faith’s phylogenetic diversity (Faith’s PD), calculated with the picante package (Kembel et al. 2010), represents the sum of branch lengths separating taxa within a community. Standardized effect sizes (SES) were calculated as z-scores based on 999 simulations and used to compare treatments while correcting for differences in richness. Among-sample dissimilarity was calculated using Bray–Curtis and weighted UniFrac distances after rarefaction to 257 sequences.

### 2.7 Statistical analyses

Measurements generated three sub-experimental datasets: thermal tolerance, gut morphology, and gut microbiome composition (Section 2.4). Within each dataset, we independently tested the effects of temperature, food quality, and their interaction. Growth and developmental rate were analyzed across all measured tadpoles, with developmental stage included as a covariate to account for minor stage differences caused by scheduling constraints. Analyses were conducted in R v4.3.1, except those of microbiome composition, which used R v4.5.1 (R Development Core Team 2021). Growth, developmental rate, thermal tolerance, and gut morphology were analyzed using comparable approaches, whereas microbiome composition required a separate workflow (Section 2.6).

#### 2.7.1 Growth, development, thermal tolerance, and gut morphology

Analyses used tidyverse for data manipulation (Wickham et al. 2019), ggplot2 for visualization (Wickham 2016), nlme (Pinheiro et al. 2024), lme4 (Bates et al. 2015), and lmerTest (Kuznetsova et al. 2017) for mixed-effects modelling, rstatix for descriptive statistics and outlier detection (Kassambara 2023), car for testing variance homogeneity (Fox & Weisberg 2019), and emmeans for estimated marginal means and post-hoc comparisons (Lenth 2026).

Separate models were fitted for growth rate, developmental rate, CT_min_, CT_max_, relative gut length, and gut mass, with temperature, food quality, and their interaction as fixed effects. Because tadpoles originated from five clutches, Clutch ID was included as a random effect in all initial models and removed only when its variance was effectively zero. Initial linear mixed-effects models were fitted with lme4::lmer(), including additional random effects and response-specific covariates where appropriate. Diagnostics included residual plots, normality tests, Levene’s tests, and assessment of random-effect variance. Final models were simplified or adjusted as indicated by these diagnostics. Statistical significance was assessed at α = 0.05.

When main effects or interactions were significant, estimated marginal means for all treatment combinations were calculated with emmeans(). Pairwise contrasts were Tukey-adjusted, and significant differences were summarized using compact letter displays. Table S3 reports EMMs, standard errors, degrees of freedom, and 95% confidence intervals for all treatment combinations. Full diagnostics and final model structures are provided in the Electronic Supplementary Material (Tables S1–S2; Figs. S1–S6).

### 2.7.2 Microbiome diversity and composition

To test whether food quality, temperature or their interaction (as fixed variables) influenced alpha diversity (Hill numbers and Faith’s PD), we built a mixed model using the package *afex* (Singmann et al. 2024), including clutch as a random variable. Post hoc tests were performed with *emmeans* (Lenth 2026).

Beta diversity based on Bray–Curtis and weighted UniFrac distances was compared across temperature, food quality, and their interaction by PERMANOVA using *adonis2* in the vegan package (Oksanen et al. 2026), with group dispersion tested using *betadisper*. A phyloseq object was created with phyloseq (McMurdie & Holmes 2013). Taxa contributing most strongly to treatment differences in microbiome composition were identified by linear discriminant analysis effect size (LEfSe; Segata et al. 2011) using microbiomeMarker (Cao et al. 2022). We retained LDA scores >4, which yielded results consistent with ALDEx2 (Eterovick et al. 2026), a probabilistic method robust to variation in read counts (Fernandes et al. 2013).

Taxa associated with both dietary nutrient assimilation (δ^¹⁵^N and δ^¹³^C) and health-related traits (body condition and developmental rate) were identified using lasso-penalized generalized linear mixed models in FLORAL (Fei et al. 2024), which detects biomarkers explaining variation in continuous variables. Because many reads could not be assigned to species, associations were tested at the genus and family levels. To assess food × temperature interactions, analyses were also conducted separately at each temperature; taxa identified as biomarkers at both temperatures were considered further evidence of food effects independent of temperature. Metabolic pathways were inferred from 16S rRNA gene data using PICRUSt2 (Douglas et al. 2020), and their differential abundance was analyzed with ALDEx2 in ggpicrust2 (Yang et al. 2023).

## 3. Results

### 3.1 Treatment effects on growth, development, thermal tolerance, and intestinal morphology

#### 3.1.1 Growth and developmental rate

Growth rate increased with temperature (*F*_₁,₃₁₂_ = 40.49, *p* < 0.001) and food quality (*F*_₂,₃₁₂_ = 7.30, *p* = 0.001), without a significant temperature × food-quality interaction (*F*_₂,₃₁₂_ = 2.03, *p* = 0.133; Fig. S7A; Table 1). Back-transformed estimated marginal means ranged from 14.4 ± 2.0 mg day⁻¹ under low food at 18 °C to 25.3 ± 3.6 mg day⁻¹ under high food at 24.5 °C (Table S3). Tukey-adjusted contrasts showed higher growth at 24.5 than at 18 °C across all food levels (*t* = 3.21–8.86, *p* ≤ 0.018). Within the 24.5 °C treatment, growth was higher under high than under low or medium food quality (low vs. high: *t* = −3.21, *p* = 0.018; medium vs. high: *t* = −3.28, *p* = 0.015).

**Table 1.**
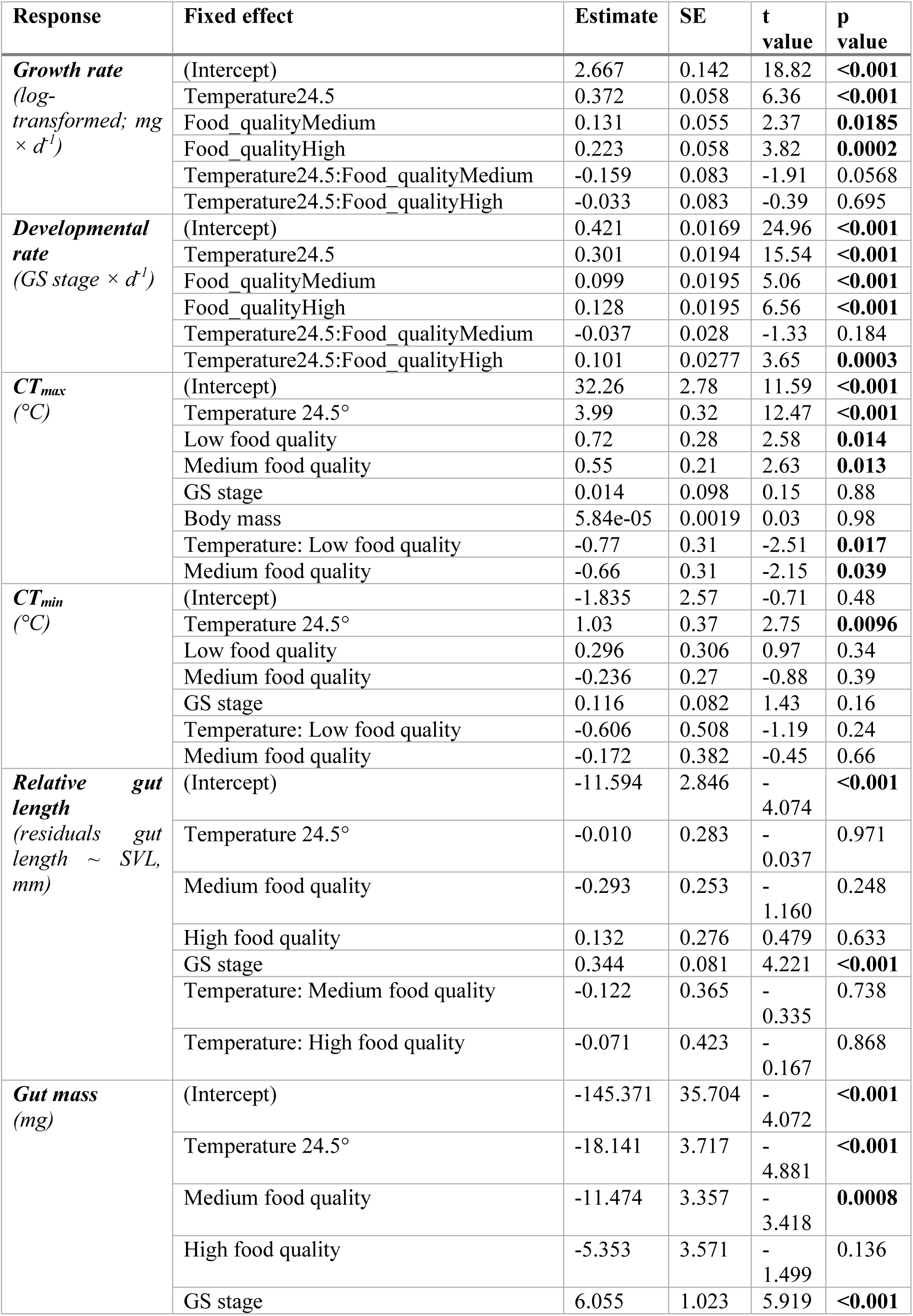
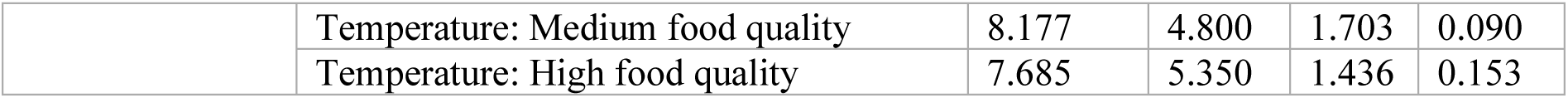
Fixed effect estimates from linear mixed-effects models (LMMs) or linear models (LMs) for six response variables in *Rana temporaria* tadpoles: growth rate, developmental rate, CT_max_, CT_min_, relative gut length, and gut mass. Shown are coefficients (Estimate), standard errors (SE), t-values, and p-values for temperature (18 °C vs. 24.5 °C), food quality (low, medium, high), Gosner stage (GS_stage), and their interactions where included. Random intercepts for Clutch or Experiment were included as appropriate. Significant effects (p < 0.05) are indicated in bold.

Developmental rate increased with temperature (*F*_₁,₃₁₂_ = 241.59, *p* < 0.001) and food quality (*F*_₂,₃₁₂_ = 23.96, *p* < 0.001), with a significant temperature × food-quality interaction (*F*_₂,₃₁₂_ = 12.88, *p* < 0.001; Fig. S7B; Table 1). Back-transformed estimated marginal means ranged from 0.421 ± 0.017 GS day⁻¹ under low food at 18 °C to 0.952 ± 0.017 GS day⁻¹ under high food at 24.5 °C (Table S3). Tukey-adjusted contrasts showed faster development at 24.5 than at 18 °C across all food levels (*t* = 3.01–27.36, *p* ≤ 0.034). Food-quality effects occurred at both temperatures but were strongest at 24.5 °C, where development was faster under high than under low or medium food quality (low vs. high: *t* = −11.61, *p* < 0.001; medium vs. high: *t* = −8.26, *p* < 0.001).

#### 3.1.2 Thermal tolerance

Thermal tolerance was strongly affected by temperature for both CT_min_ and CT_max_.

CT_min_ increased with temperature (*F*_₁,₃₃_ = 31.62, *p* < 0.001), whereas neither food quality (*F*_₂,₃₃_ = 1.89, *p* = 0.167) nor the temperature × food-quality interaction was significant (*F*_₂,₃₃_ = 0.71, *p* = 0.498; Fig. S7D; Table 1). Estimated marginal means were lower at 18 °C than at 24.5 °C across all food treatments (Table S3). Tukey-adjusted contrasts detected no differences among food treatments within temperatures (*t* ≤ 1.88, *p* ≥ 0.16) or between temperatures within individual food treatments (*t* ≤ 1.68, *p* ≥ 0.10).

CT_max_ increased strongly with temperature (*F*_₁,₃₅_ = 164.87, *p* < 0.001) and showed a significant temperature × food-quality interaction (*F*_₂,₃₅_ = 3.87, *p* = 0.030), whereas the main effect of food quality was not significant (*F*₂,₃₃ = 1.63, *p* = 0.210; Fig. S7C; Table 1). Estimated marginal means ranged from 32.9–33.6 °C at 18 °C to 36.5–36.6 °C at 24.5 °C (Table S3). Tukey-adjusted contrasts showed higher CT_max_ at 24.5 than at 18 °C within each food treatment (*t* = 8.44–12.36, *p* < 0.001). Food-quality effects were weak and detected only at 18 °C, where CTmax was slightly lower under high than low food quality (*t* = −2.47, *p* = 0.047).

#### 3.1.3 Intestinal morphology

Relative gut length was affected by food quality (*F*_₂,₁₄₀_ = 5.27, *p* = 0.006) and developmental stage (*F*_₁,₁₄₀_ = 25.62, *p* < 0.001), but not by temperature (*F*_₁,₁₄₀_ = 3.34, *p* = 0.070) or the temperature × food-quality interaction (*F*_₂,₁₄₀_ = 0.06, *p* = 0.945; Fig. S7E; Table 1). Estimated marginal means were highest under high food quality (18 °C: 0.22 ± 0.19; 24.5 °C: 0.14 ± 0.19) and lowest under medium food quality (18 °C: −0.21 ± 0.19; 24.5 °C: −0.34 ± 0.20; Table S3). Relative gut length showed no consistent differences between temperatures within food treatments and increased with developmental stage.

Gut mass was affected by temperature (*F*_₁,₁₆₂_ = 74.12, *p* < 0.001), food quality (*F*_₂,₁₆₂_ = 8.54, *p* = 0.0003), and developmental stage (*F*_₁,₁₆₂_ = 55.75, *p* < 0.001), without a significant temperature × food-quality interaction (*F*_₂,₁₆₂_ = 1.72, *p* = 0.183; Fig. S7F; Table 1). Estimated marginal means ranged from 60.1 ± 2.47 mg under low food quality at 18 °C to 38.7 ± 2.57 mg under medium food quality at 24.5 °C and were consistently lower at 24.5 than at 18 °C across food treatments (Table S3). Gut mass also increased with developmental stage.

### 3.2 Microbiome composition, biomarkers, and predicted effects across treatments

Bacterial community diversity (Hill numbers) was not influenced by temperature (df = 1, 44.306; F = 2.972; p = 0.092), food treatment (df = 2, 45.036; F = 1.021; p = 0.369), or their interaction (df = 2, 45.663; F = 2.472; p = 0.096; Fig. S7A). However, phylogenetic diversity was influenced by food treatment (df = 2, 45.972; F = 7.635; p = 0.001), temperature (df = 1, 45.353; F = 13.739; p = 0.001), and their interaction (df = 2, 46.250; F = 3.762, p = 0.031). It was lower at low and medium food quality at 18 °C compared to high food quality at 18 °C and all treatments at 24.5 °C (Fig. S9B).

Microbiome composition based on Bray-Curtis dissimilarity estimates differed among food treatments (pseudo-F = 3.78, n = 53, p = 0.001) and between temperatures (pseudo-F = 17.04, n = 53, p = 0.001), and there was an interaction between the two factors (pseudo-F = 3.17, n = 53, p = 0.001)(Fig. 1A). At 18 °C, all food treatments were different from one another (pairwise comparisons all p < 0.009). At 24.5 °C, the low and high food quality treatments were different from one another (pairwise comparison: Low-High p = 0.015), but neither differed from the medium food treatment (pairwise comparisons: Low-Med p = 0.07, Med-High p = 0.567). Results were similar for weighted Unifrac distances: beta-diversity estimates again differed among food treatments (pseudo-F = 2.51, n = 53, p = 0.003) and strongly between temperatures (pseudo-F = 10.05, n = 53, p = 0.001), and their interaction (pseudo-F = 3.10, n = 53, p = 0.003). In pairwise comparisons, all food treatments differed from one another at 18 °C (all comparisons p < 0.027), but did not at 24.5 °C (all p > 0.103). Group dispersion did not differ among treatment combinations (temperature, diet) for either dissimilarity metric (Bray-Curtis p = 0.89, Unifrac p = .80).

**Fig. 1.**
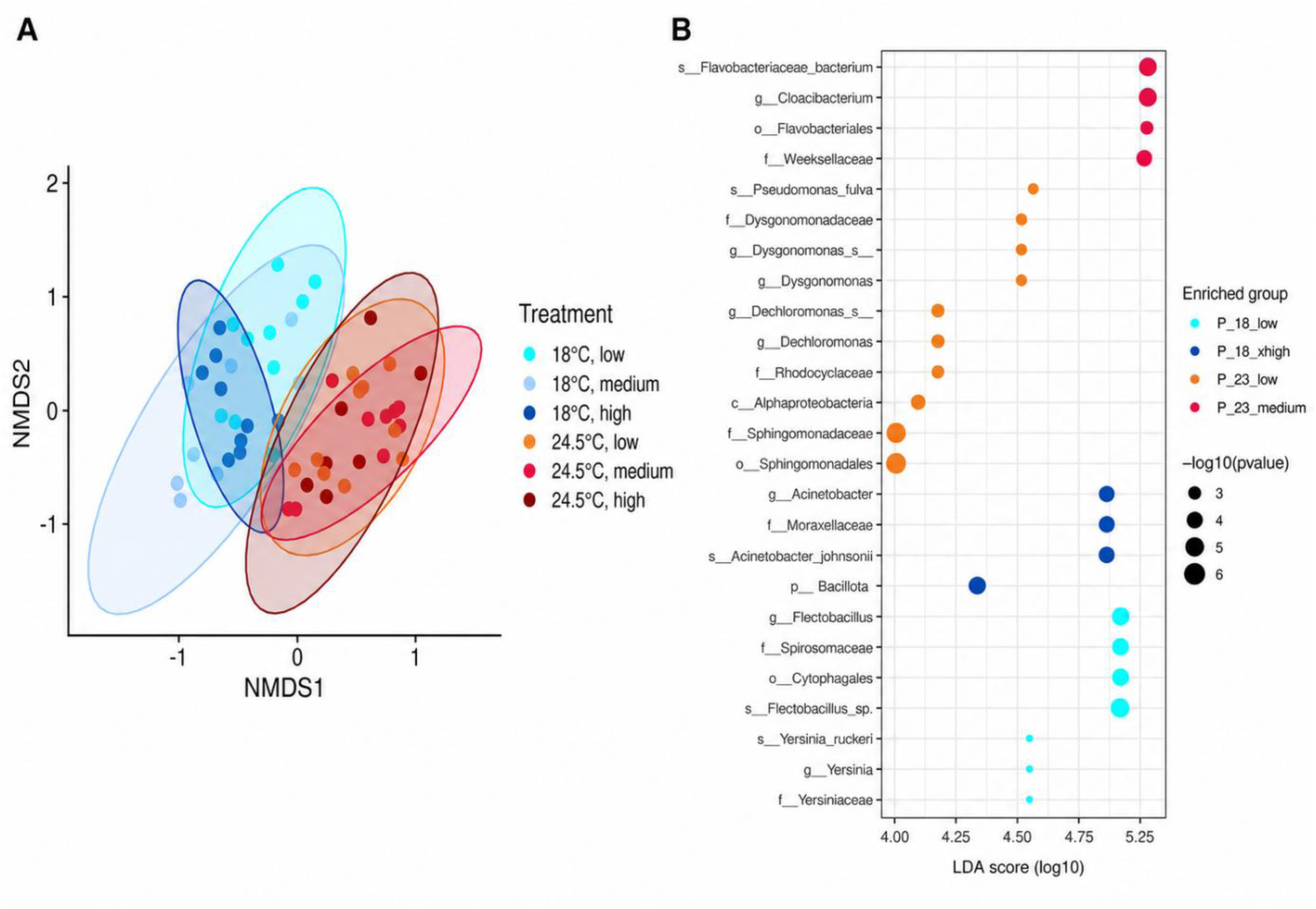
Variation in bacterial community composition and structure in the guts of larvae of *Rana temporaria* reared at different temperatures and food quality (A) and taxonomic groups identified as microbiome biomarkers with significant increase within specific treatments (B). Ordination in (A) was constructed using non-metric multidimensional scaling based on Bray- Curtis dissimilarity (stress = 0.186).

Larvae reared at 18 °C and low food quality had *Flectobacillus* (Spirosomataceae, Cytophagales) and *Yersinia ruckeri* (Yersiniaceae), whereas larvae reared at 18 °C and high food quality had *Acinetobacter johnsonii* (Moraxellaceae) and an unclassified Bacillota as indicator microbial taxa (Fig. 2B, Fig. S10). Larvae reared at 24.5 °C and low food quality had *Pseudomonas fulva, Dysgonomonas* (Dysgonomonadaceae), *Dechloromonas* (Rhodocyclaceae) and an unclassified Sphingomonadaceae (Sphingomonadales, Alphaproteobacteria), whereas larvae reared at 24.5 °C and medium food quality had *Cloacibacterium* (Flavobacteriales, Weeksellaceae) as indicator microbial taxa (Fig. 1B, Fig. S10).

**Fig. 2.**
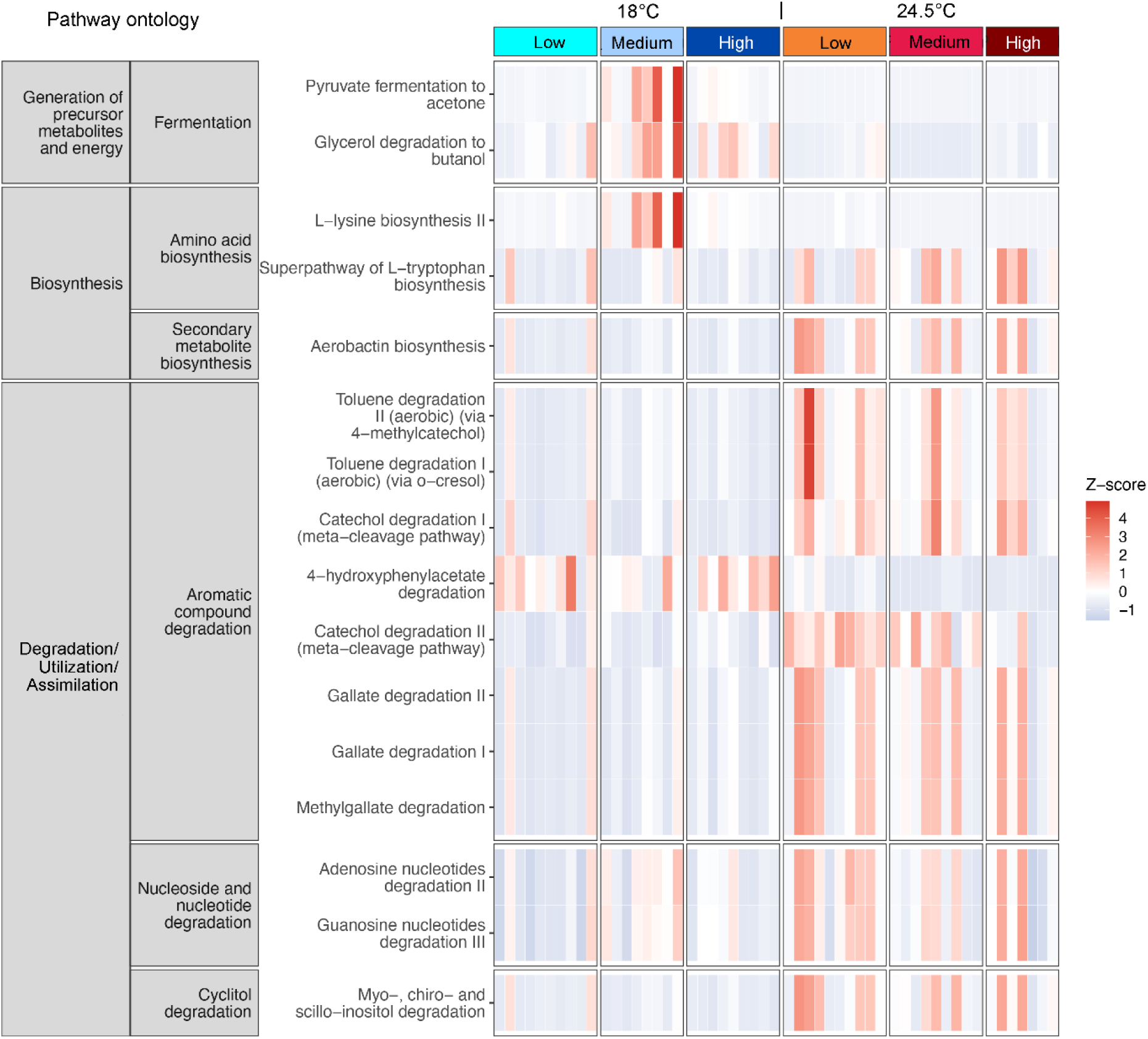
Heatmap showing predicted expression of the 15 top significant pathways by the gut bacteria of *Rana temporaria* larvae across treatments crossing two rearing temperatures and three food regimes. Higher z-scores (shaded red) indicate enrichment of a given functional pathway in the sample; lower z-scores (shaded blue) indicate the pathway is depleted.

*Flectobacillus* (Spirosomaceae) was more abundant at 18 °C and negatively associated with food quality, linking it to slower growth. Reads assigned to *Acetinobacter* or Yersiniaceae were negatively associated with developmental rate but not clearly with food assimilation, suggesting a primarily temperature-dependent response. At 24.5 °C, *Pseudomonas* (Pseudomonadaceae), *Dysgonomonas* (Dysgonomonadaceae), and *Novosphingobium* (Sphingomonadaceae) were negatively associated with food quality. The SILVA-designated genera env.OPS_17 (Sphingomonadales) and NS11-12_marine_group (Sphingobacteriales) were positively associated with nitrogen and carbon assimilation at the higher temperature (Glöckner et al. 2017; Quast et al. 2012; Yilmaz et al. 2014; Fig. S8).

The 15 most significant predicted metabolic pathways differed primarily between temperatures, although most also interacted with food quality. Fermentation and L-lysine biosynthesis pathways generally increased at 18 °C under medium food quality (Fig. 2; Figs. S11–S12). In contrast, degradation, utilization, and assimilation pathways generally increased at 24.5 °C across food treatments, except 4-hydroxyphenylacetate degradation, which increased at 18 °C. At low food quality, warming increased pathways for guanosine (PWY-6608), adenosine (SALVADEHYPOX-PWY), gallate, and methylgallate degradation (GALLATE- DEGRADATION-I and -II; METHYLGALLATE-DEGRADATION-PWY). Catechol degradation I (PWY-5415) increased with temperature under low and high food quality, whereas catechol degradation II (PWY-5420) increased under low and medium food quality. Toluene degradation pathways PWY-5180 and PWY-5182 increased under low and high food quality, respectively, while myo-, chiro-, and scyllo-inositol degradation (PWY-7237) increased with temperature irrespective of food quality. Aerobactin biosynthesis also increased with temperature, particularly under low and high food quality.

Pathways with increased expression at low rearing temperatures included 4- hydroxyphenylacetate degradation at low or high food quality, pyruvate fermentation to acetone (PWY-6588), especially at medium food quality, glycerol degradation to butanol (PWY-7003) and L-lysine biosynthesis II (PWY-2941) at medium food quality (Fig. S12).

## 4. Discussion

### 4.1 Growth, development, and thermal tolerance

Consistent with established patterns in ectotherms, warming accelerates larval growth and development (Ruthsatz et al. 2018; Sinai et al. 2022; Albecker et al. 2023) and increases upper thermal limits via acclimation (Pottier et al. 2022; Ruthsatz et al. 2022a, 2024). Accordingly, larvae reared at 24.5 °C grew and developed faster and exhibited higher CT_max_ than those reared at 18 °C. However, these temperature-driven plastic responses are energetically costly: when exposed to higher temperatures, metabolic rate and baseline maintenance costs increase, and sustaining rapid growth and development requires a sufficient surplus of assimilable energy above those maintenance demands. At the same time, both thermal acclimation and tolerance to acute heat exposure are likely to entail additional energetic and biochemical investment (e.g., cellular protection and repair; Feder & Hofmann 1999; Somero 2020), meaning that expressing higher CT_max_ under warming should be easiest when energy supply is ample (Hardison & Eliason 2024). Larvae in the current study exhibited these heat tolerance plastic responses at a gradient of food quality while also increasing growth and development. The observed differences among food treatments show that energy supply (i.e., food quality) matters, but the response of the gut bacterial community to the available nutrient sources likely contributed to deployment of compensatory strategies by the larvae. The lack of a trade-off between growth, development, and heat tolerance indicate maintenance of suitable net energy gain under warming, with digestive adjustments, whether host- or microbially-mediated, being a plausible route.

### 4.2 Gut morphology

Phenotypic flexibility of the digestive tract can help vertebrates maintain energy gain when resource quality declines or energetic demand rises (Secor 2001). Animals may rapidly adjust gut length, diameter, surface structure, and tissue investment to increase processing of low-quality food or reduce the maintenance costs of digestive tissue when intake is limited (Yang & Joern 1994; Piersma & Lindström 1997; Secor 2001). In amphibian larvae, poor-quality diets often induce gut elongation, potentially improving nutrient extraction from fibrous or nutrient-poor resources (Álvarez & Nicieza 2002; Ruthsatz et al. 2019, 2022b).

In the present study, however, morphological compensation was limited. Relative gut length varied only weakly with food quality, was unaffected by temperature, and increased with Gosner stage, consistent with ontogenetic scaling (Hourdry et al. 1996). By contrast, gut mass declined at higher temperature and under low-quality diets. This contrasts with previous studies in which poor-quality diets induced longer intestines, including leaf-litter diets and diets with reduced protein and nutrient density (Stoler & Relyea 2013; Ruthsatz et al. 2019). Together, these discrepancies suggest that gut elongation is not a universal response to energetic constraint but may depend on the nutritional contrast, the developmental window sampled, and/or the presence of concurrent stressors such as elevated temperature.

A methodological caveat is that guts were not flushed prior to weighing to preserve tissue integrity for length measurements, and gut-content density could vary among diets; however, this is unlikely to explain the strong and consistent temperature effect on gut mass. More broadly, reduced gut mass under warming argues against a simple “*build more gut under stress*” model and may instead reflect allocation toward rapid development and heat-tolerance acclimation, or constraints on maintaining costly gut tissue under poor diets (Secor 2001). Larvae may consequently rely more on less costly responses, including microbial community shifts, to sustain digestion under thermal and nutritional stress.

### 4.3 Gut microbiome

The gut microbiome is increasingly recognized as an environmentally responsive component of host phenotype that influences nutrient acquisition, metabolism, and stress tolerance (Youngblut et al. 2019; Sepulveda & Moeller 2020). Because gut microbiomes respond faster to environmental changes than their hosts (Voolstra & Ziegler 2020), microbial shifts may reduce the need for energetic investment in host digestive remodeling, as may have occurred in our study. Indeed, both diet and temperature have been shown to restructure gut bacterial communities in ectotherms, with consequences for performance and fitness (Greenspan et al. 2020; Suzzi et al. 2023; Zhu et al. 2024). Furthermore, experiments using amphibian larvae have shown that microbiome disruption can reduce thermal tolerance and survival under heat stress, suggesting that microbiota have a beneficial role in modulating responses to extreme temperatures (Fontaine et al. 2022). Nevertheless, studies integrating temperature–diet interactions remain scarce (but see Fontaine et al. 2018; rev. in Hardison & Eliason 2024), making our combined analysis of both stressors and host performance particularly relevant for climate change contexts.

Larvae of *R. temporaria* showed the strongest compositional shifts under high temperature combined with low diet quality, representing a dual constraint of elevated metabolic demand and limited energy assimilation. Under these conditions, indicator taxa included metabolically versatile lineages such as *Pseudomonas*, *Dysgonomonas*, *Sphingomonadaceae*, *Sphingomonadales* (including env.OPS_17), and *Dechloromonas*. These taxa are consistent with a community adapted to flexible substrate use and chemical processing, and potentially to mitigating heat-induced oxidative stress. For example, *Pseudomonas* and *Sphingomonas* possess pathways for degrading gallate, methylgallate, and catechol (Karp et al. 2019), compounds with antioxidant properties found in plants (Woodman 2009). At 24.5 °C, *Pseudomonas* and *Sphingomonadaceae* were negatively correlated with food quality, suggesting increased degradation of such compounds under low-quality diets. Admittedly, linking microbial shifts to host phenotypes requires caution; these changes in the microbiota may or may not benefit the host (Sepulveda & Moeller 2020). Our results, however, support such associations, as individuals reared at elevated temperatures showed faster development, increased growth, and higher CT_max_ despite lower food quality. This aligns with evidence that microbiomes can influence host thermal biology (Fontaine et al. 2022) and that microbiome manipulation can predictably alter ectotherm heat tolerance (Dallas et al. 2024).

Although 16S-based functional predictions reflect potential rather than actual gene expression, they are valuable for hypothesis generation (Caspi et al. 2016). Under elevated temperature, particularly with low food quality, predicted pathways converged on three key functions: (1) expanded carbon acquisition from complex substrates, (2) resource economy via nucleotide recycling, and (3) micronutrient acquisition and stress resilience, particularly through iron scavenging. Pathways such as gallate, catechol, toluene, and inositol degradation suggest an enhanced capacity to utilize diverse carbon sources and process plant-derived compounds (Caspi et al. 2016). This is relevant because low-quality plant diets are rich in phenolic compounds with antioxidant properties (e.g., barley; Hajji et al. 2018). Increased purine and nucleoside salvage pathways at higher temperatures suggest enhanced resource recycling under nutrient limitation, potentially stabilizing microbial function and host-associated benefits. Additionally, increased aerobactin biosynthesis indicates enhanced iron acquisition, which can support bacterial growth and mitigate oxidative stress (Li et al. 2021). Aerobactins bind ferric ions, facilitating iron uptake by bacteria (Karp et al. 2019). As iron is often limiting, increased aerobactin production likely promotes microbial growth and associated metabolic activity. Genera such as *Yersinia* and *Klebsiella* encode this pathway; *Yersinia* was associated with low temperature and low food quality, whereas *Klebsiella* was more abundant at higher temperatures and food quality, correlating with developmental rate. The increase of this pathway across treatments suggests that different taxa contribute to similar functional outcomes, reflecting functional redundancy in microbial communities (Louca et al. 2018).

### 4.4. Conclusions

In this study, we show that amphibian larvae can maintain substantial performance under a climate-relevant combination of stressors through strong plasticity in growth and developmental rate and by acclimating heat tolerance to warmer conditions. Host gut morphology changed only modestly, whereas the gut microbiome showed the clearest plastic response under energetic constraint, with the greatest compositional turnover and predicted functional shifts occurring when warming coincided with low food quality. Our findings add to growing evidence that microbiota can influence host thermal biology (Hardison & Eliason 2024). Nevertheless, we did not test causality directly, and our functional inferences are based on 16S-derived predictions of metabolic capacity rather than measured pathway expression or metabolite production. Validating the proposed mechanisms will require metagenomics/metatranscriptomics and/or metabolomics under warming and low food quality and direct causality tests using microbiome depletion, transplants, or gnotobiotic approaches, as already shown feasible and informative in tadpoles (Fontaine et al. 2022; Dallas et al. 2024). Because gut microbiomes can shape host nutrition, immunity, and stress tolerance, and thus potentially influence population persistence under global-change-driven environmental stress, mechanistic and causal tests of microbial buffering should become a more explicit component of global-change biology and conservation. We therefore encourage additional research on the role of microbiomes in host ability to deal with environmental stressors, and on the mechanisms involved, to enable conservation strategies that account for holobiont adaptive capacity.

## Acknowledgements

We thank Miguel Vences, Sven Gippner, and Janina Rudolph for field assistance, and Lara Keunecke, Fabian Bartels, and Ben Oetken for assistance in animal husbandry. We are grateful to Arne Wulff for his aid in the bomb calorimetry at the University of Hamburg.

## Author contributions

KR: conceptualization, funding acquisition, project administration, supervision, methodology, investigation, formal analysis, data curation, visualization, writing: original draft, writing: review and editing

MCH: methodology, formal analysis, writing: review and editing

MT: investigation, data curation, writing: review and editing

MdA: investigation, data curation, writing: review and editing

JG: investigation, writing: review and editing

PCE: conceptualization, methodology, investigation, formal analysis, data curation, visualization, writing: original draft, writing: review and editing

## Statement of Ethics

The authors have no ethical conflicts to disclose. The experiments were conducted under permission from the *Niedersächsisches Landesamt für Verbraucherschutz und Lebensmittelsicherheit*, Germany (Gz. 33.19-42502-04-20/3590 and 33.19-42502-04-22- 00274). Fieldwork in Lower Saxony was carried out with permits of Stadt Braunschweig (Stadt Braunschweig - Fachbereich Umwelt und Naturschutz, Willy-Brandt-Platz 13, 38102 Braunschweig; Gz. 68.11-11.8-3.3).

## Data availability

Data are deposited in Figshare (https://figshare.com/s/db8d34e0515e24cb26de) and will be published after acceptance (doi: 10.6084/m9.figshare.33095144). Raw sequences are deposited in the NCBI (BioProject PRJNA1304763).

## Statement of inclusion

Our study was conducted in Germany, with field sampling and experimental work based in the region where the study organisms were collected. The research involved scientists with relevant regional and international expertise, who contributed to the development, analysis, interpretation, and communication of the research. Whenever relevant, we considered and cited research conducted on the study species and related ecological questions within the region.

