## Supplementary Material for "Microbiome plasticity, not gut morphology, is linked to amphibian larval performance under elevated temperatures and low food quality"

### Statistical analysis

*A. Growth, development, thermal tolerance, and gut morphology*

#### a. General diagnostic and preprocessing procedures

Prior to analyses, data were screened for completeness and extreme outliers using the *identify_outliers()* function from rstatix. Model assumptions of normality and homoscedasticity were assessed for all dependent variables using standard diagnostic plots (histograms, Q–Q plots, residual vs. fitted, boxplots, scale–location, and Cook's distance/leverage plots; Fig. S1–S6) as well as Shapiro–Wilk and Levene's tests (Table S1).

All diagnostic statistics reported refer to the initial models (i.e., null models), which included relevant random effects and covariates. Based on these diagnostics, final models (Table S2) were fitted with simplifications (e.g., removal of random effects or covariates) or transformations applied where necessary to improve normality, stabilize variance, or enhance model interpretability. Full details of model diagnostics, simplifications, and final model structures are provided in Tables S1–S2 and Figures S1–S6.

#### b. Trait-specific analytical notes

**Growth and developmental rate** – Data screening identified two missing/outlier values, which were removed (<1% of cases).

> **Growth rate:** Initial diagnostics on the LMM with temperature, food quality, and their interaction as fixed effects, and random intercepts for 'Experiment' and 'Clutch ID' revealed strong right skew and heteroscedasticity across temperature × food combinations. Log-transformation improved normality, but residual variance heterogeneity persisted (Shapiro-Wilk: W=0.955, p<0.001; Levene's test: F= 6.96, p<0.001; Table S1; Fig. S1). 'Experiment' was retained as a random intercept, whereas 'Clutch ID' contributed negligible variance and was removed. The final model was fitted as a weighted LMM using the *lme()* function in the nmle package with a varIdent variance structure allowing separate residual variances for each temperature × food combination (weights = varIdent(form = ~1 | Temp:Food); Table S2). Fixed effects included temperature, food quality, and their interaction.
>
> **Developmental rate:** Initial model diagnostics showed approximate symmetry and weak heteroscedasticity (Shapiro-Wilk: W=0.968, p<0.001; Levene's test: F= 12.95, p<0.001; Table S1; Fig. S2). No transformation or variance weighting was required. 'Experiment' was retained as a random intercept for consistency, whereas 'Clutch ID' contributed no detectable variance and was removed. The final unweighted LMM model included temperature, food quality, and their interaction as fixed effects and was fitted using the *lme()* function in the nlme package, following the same structure as the growth rate model (Table S2)."

**Thermal tolerance ($CT_{max}$ and $CT_{min}$)** – 'Gosner stage (GS stage)' and 'body mass' were included to account for size- and stage-dependent variation, and 'initial measurement time' to account for potential diurnal effects. Collinearity was assessed using VIF. No outliers required removal.

**$CT_{max}$**: Initial diagnostics of the LMM with temperature, food quality, and their interaction as fixed effects, Clutch ID as a random intercept, and covariates GS stage, body mass, and initial time revealed residual non-normality due to a multimodal distribution (~32–33°C, ~35°C, ~36–37°C; Shapiro-Wilk: W=0.85, p<0.001) but approximately homogeneous variance across treatments (Levene's test: F= 2.34, p=0.057; Table S1; Fig. S3). Given the robustness of LMMs to moderate non-normality, untransformed $CT_{max}$ values were retained. Clutch-level variance was small but retained to account for sibling non-independence. All covariates had VIF < 5 (GS stage: 2.07; body mass: 1.68; initial measurement time: 1.06). The final model retained all fixed effects and covariates in the initial LMM structure using the *lmer()* function in the lme4 package.

**$CT_{min}$**: Initial LMM diagnostics indicated mild non-normality (Shapiro-Wilk: W=0.94, p=0.014) and homoscedasticity (Levene's test: F=1.24, p=0.31; Table S1; Fig. S4). The random effect for Clutch ID was singular (variance = 0), and GS stage and body mass were strongly collinear (r=0.837; GVIFs < 5 but elevated; GS stage: 6.1; body mass: 3.72; initial measurement time: 2.97). To improve model stability, Clutch ID was removed, and body mass was excluded due to collinearity, while GS stage and initial measurement time were retained along with temperature, food quality, and their interaction as fixed effects. The simplified linear model showed acceptable residual structure, low collinearity (GVIFs < 2), and homoscedasticity, providing a biologically defensible framework (Table S2).

**Relative gut length and gut mass** – All intestinal measures are morphometric traits that typically scale with body size (Relyea & Auld 2004). To account for body size effects, we calculated relative gut length using residuals from linear regressions of gut length on snout–vent length (SVL) across the full dataset (i.e., all treatment groups; Ruthsatz et al. 2019, 2022) prior to statistical analysis. Positive residuals indicate individuals with relatively long intestines for their body size, whereas negative residuals indicate relatively short intestines. These residuals were used as the response variable in all subsequent statistical analyses.

Developmental stage (GS stage) was included as a covariate to account for ontogenetic changes in intestinal morphology (Shi 2000).

Initial LMM diagnostics with Clutch ID as a random effect showed no extreme outliers and acceptable normality (Shapiro-Wilk – relative gut length: W=0.99, p=0.413; gut mass: W=0.98, p=0.034) and variance homogeneity homogeneity (Levene's test – relative gut length: F=1.69, p=0.138; gut mass: F=1.84, p=0.107; Table S1; Fig. S5–S6). In both traits, the random effect of Clutch ID had zero variance and was removed to improve model stability. Developmental stage (GS stage) was retained as a covariate. Final analyses used standard linear models with temperature, food quality, and their interaction as fixed effects, and GS stage as a covariate. Residuals confirmed acceptable normality and homoscedasticity (Table S2).

c. **Diagnostic summary table (Table S1)**

**Table S1.** Outlier screening, assumption tests, data transformations, random-effect retention, and final diagnostic outcomes for each dependent variable. Relative gut length was calculated as residuals from regressions of gut length on snout-vent-length (SVL) across all treatment groups. GS stage = developmental stage *sensu* Gosner (1960). Initial measurement time= Start of CT measurements. LM = linear model. LMM = linear mixed-effects model.

| Dependent Variable | Outliers Removed | Transformation | Random Effects (final) | Shapiro-Wilk (W, p) | Levene's Test (F, p) | Covariates / Notes |
|---|---|---|---|---|---|---|
| **Growth rate (mg × $d^{-1}$)** | 2 | log | Experiment | $W$=0.955, $p$<0.001 | $F$=2.28, $p$=0.0634 | Weighted LMM (varIdent) to address heteroscedasticity; Clutch removed |
| **Developmental rate (GS × $d^{-1}$)** | 2 | none | Experiment | $W$=0.968, $p$<0.001 | $F$=12.95, $p$<0.001 | Unweighted LMM; Clutch ID removed due to zero variance |
| **$CT_{max}$ (°C)** | 0 | none | Clutch ID | $W$=0.850, p<0.001 | $F$=2.34, $p$=0.057 | LMM retained as initial structure; covariates: GS stage, body mass, initial measurement time |
| **$CT_{min}$ (°C)** | 0 | none | None | $W$=0.940, $p$=0.014 | $F$=1.24, $p$=0.310 | LM fitted after removing Clutch ID and body mass due to singularity and collinearity; covariates: GS stage and initial measurement time |
| **Relative gut length** | 0 | none | None | $W$=0.990, $p$=0.413 | $F$=1.69, $p$=0.138 | LM; covariate: GS stage; Clutch ID removed due to zero variance |
| **Gut mass (mg)** | 0 | none | W= | $W$=0.980, $p$=0.034 | $F$=1.84, $p$=0.107 | LM; covariate: GS stage; Clutch ID removed due to zero variance |

### d. Final model specifications (Table S2)

**Table S2.** Final model structures for each response variable, including formula notation, fixed and random effects, transformations, weighting, and the modelling function used.

| Dependent Variable | Final Model Formula | Fixed Effects | Random Effects | Transformation | Weighting | Modelling Function (package:function) |
|---|---|---|---|---|---|---|
| **Growth rate (mg × d$^{-1}$)** | log(growth_rate) ~ Temperature * Food quality | Temperature, Food quality, Temp × Food | Experiment | log | varIdent(~1 | Temp:Food) |
| **Developmental rate (mg × d$^{-1}$)** | dev_rate ~ Temperature * Food quality | Temperature, Food quality, Temp × Food | Experiment | none | none | nlme::lme() |
| **$CT_{max}$ (°C)** | $CT_{max}$ ~ Temperature * Food quality + GS_stage + Mass + initial measurement time | Temperature, Food quality, Temp × Food, GS_stage, Mass, Initial_Time | Clutch ID | none | none | lme4::lmer() |
| **$CT_{min}$(°C)** | $CT_{min}$ ~ Temperature * Food quality + GS_stage + initial measurement time | Temperature, Food quality, Temp × Food, GS_stage, Initial_Time | none | none | none | stats::lm() |
| **Relative gut length** | RGL ~ Temperature * Food quality + GS_stage | Temperature, Food quality, Temp × Food, GS_stage | none | none | none | stats::lm() |
| **Gut mass (mg)** | gut_mass ~ Temperature * Food quality + GS_stage | Temperature, Food quality, Temp × Food, GS_stage | none | none | none | stats::lm() |

### e. Estimated marginal means (Table S3)

**Table S3.** Estimated marginal means (EMMs) for each temperature × food-quality treatment combination for six response variables: growth rate (GR; mg day$^{-1}$), developmental rate (DR; Gosner stages day$^{-1}$), critical thermal maximum (CTmax; °C), critical thermal minimum (CTmin; °C), relative gut length, and gut mass (mg). Values are model-predicted means (EMMs) ± standard error (SE), with associated degrees of freedom (df) and 95% confidence limits (CIL and CIU). Means are averaged over other model covariates where applicable. Temperature and food-quality combinations correspond to the experimental treatments.

| Response variable | Temperature | Food quality | emmean | SE | df | CIL | CIU |
|---|---|---|---|---|---|---|---|
| ***Growth rate (mg/day)*** | 18 | Low | 14.4 | 2.04 | 2 | 5.62 | 23.2 |
| | 24.5 | Low | 20.9 | 2.92 | 2 | 8.31 | 33.4 |
| | 18 | Medium | 16.4 | 2.28 | 2 | 6.62 | 26.2 |
| | 24.5 | Medium | 20.3 | 2.91 | 2 | 7.81 | 32.8 |
| | 18 | High | 18.0 | 2.52 | 2 | 7.16 | 28.8 |
| | 24.5 | High | 25.3 | 3.59 | 2 | 9.81 | 40.7 |
| ***Developmental rate (GS/d)*** | 18 | Low | 0.421 | 0.0169 | 2 | 0.349 | 0.494 |
| | 24.5 | Low | 0.722 | 0.0172 | 2 | 0.648 | 0.797 |
| | 18 | Medium | 0.520 | 0.0174 | 2 | 0.445 | 0.595 |
| | 24.5 | Medium | 0.784 | 0.0180 | 2 | 0.706 | 0.861 |
| | 18 | High | 0.549 | 0.0174 | 2 | 0.474 | 0.624 |
| | 24.5 | High | 0.952 | 0.0173 | 2 | 0.877 | 1.026 |
| ***CTmax (°C)*** | 18 | High | 32.9 | 0.176 | 32.3 | 32.4 | 33.4 |
| | 18 | Medium | 33.5 | 0.131 | 22.7 | 33.1 | 33.8 |
| | 18 | Low | 33.6 | 0.247 | 34.1 | 32.9 | 34.3 |
| | 24.5 | Medium | 36.5 | 0.218 | 34.4 | 35.9 | 37.1 |
| | 24.5 | Low | 36.6 | 0.171 | 26.6 | 36.1 | 37.1 |
| | 24.5 | High | 36.6 | 0.245 | 34.5 | 36.0 | 37.3 |
| ***CTmin (°C)*** | 18 | Low | 7.67 | 0.25 | 140 | 7.17 | 8.17 |
| | 24.5 | Low | 8.92 | 0.28 | 140 | 8.36 | 9.48 |
| | 18 | Medium | 7.85 | 0.27 | 140 | 7.31 | 8.39 |
| | 24.5 | Medium | 9.01 | 0.26 | 140 | 8.49 | 9.53 |
| | 18 | High | 7.90 | 0.26 | 140 | 7.38 | 8.42 |
| | 24.5 | High | 9.15 | 0.27 | 140 | 8.61 | 9.69 |
| ***Relative gut length*** | 18 | Low | 0.0882 | 0.185 | 140 | -0.277 | 0.454 |
| | 24.5 | Low | 0.0777 | 0.197 | 140 | -0.314 | 0.468 |
| | 18 | Medium | -0.2052 | 0.187 | 140 | -0.574 | 0.164 |
| | 24.5 | Medium | -0.3379 | 0.197 | 140 | -0.728 | 0.052 |
| | 18 | High | 0.2204 | 0.187 | 140 | -0.148 | 0.589 |
| | 24.5 | High | 0.1393 | 0.190 | 140 | -0.235 | 0.514 |
| ***Gut mass (mg)*** | 18 | Low | 60.1 | 2.47 | 162 | 53.5 | 66.7 |
| | 24.5 | Low | 42.0 | 2.55 | 162 | 35.2 | 48.7 |
| | 18 | Medium | 48.6 | 2.47 | 162 | 42.0 | 55.2 |
| | 24.5 | Medium | 38.7 | 2.57 | 162 | 31.8 | 45.5 |
| | 18 | High | 54.7 | 2.39 | 162 | 48.4 | 61.1 |
| | 24.5 | High | 44.3 | 2.34 | 162 | 38.0 | 50.5 |

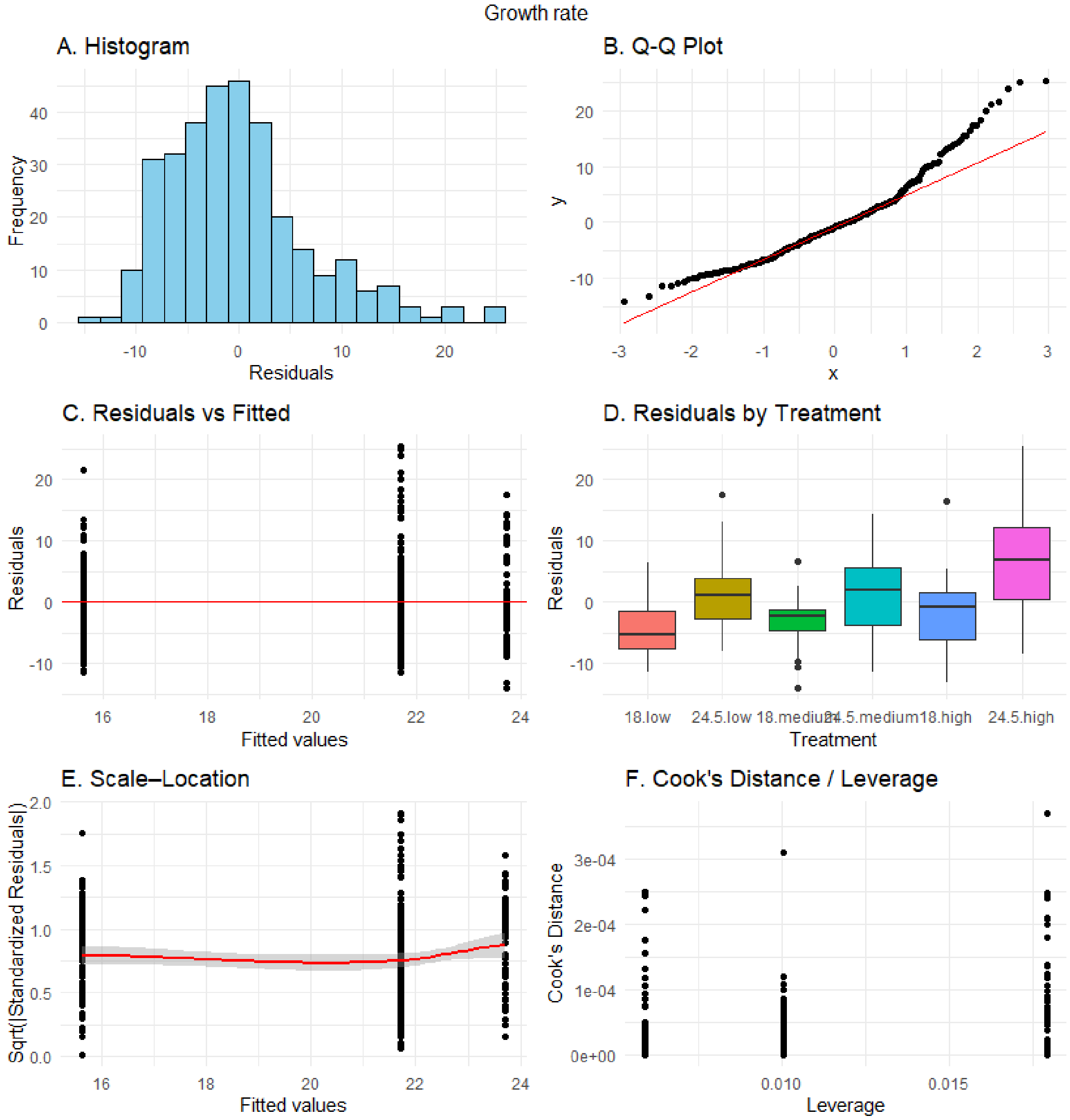


**Figure S1**. Null model assumptions of normality and homoscedasticity for growth rate (mg × $d^{-1}$) were evaluated using standard diagnostic plots: (A) histogram of residuals, (B) Q–Q plot, (C) residuals vs. fitted values, (D) residuals by treatment (Temperature × Food_quality), (E) scale–location plot, and (F) Cook's distance versus leverage plot.

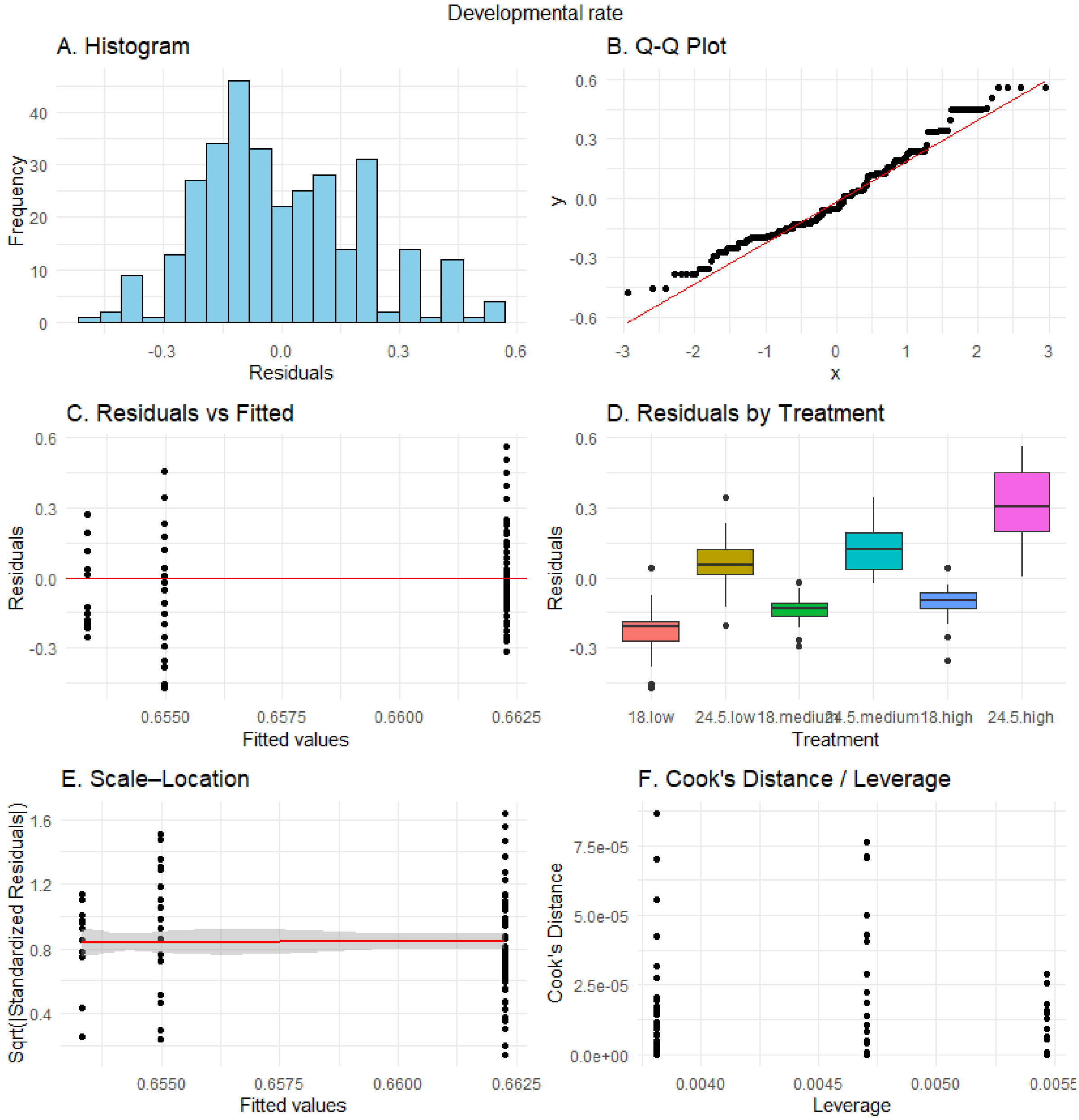


**Figure S2**. Null model assumptions of normality and homoscedasticity for developmental rate (Gosner stage × $d^{-1}$) were evaluated using standard diagnostic plots: (A) histogram of residuals, (B) Q–Q plot, (C) residuals vs. fitted values, (D) residuals by treatment (Temperature × Food_quality), (E) scale–location plot, and (F) Cook's distance versus leverage plot.

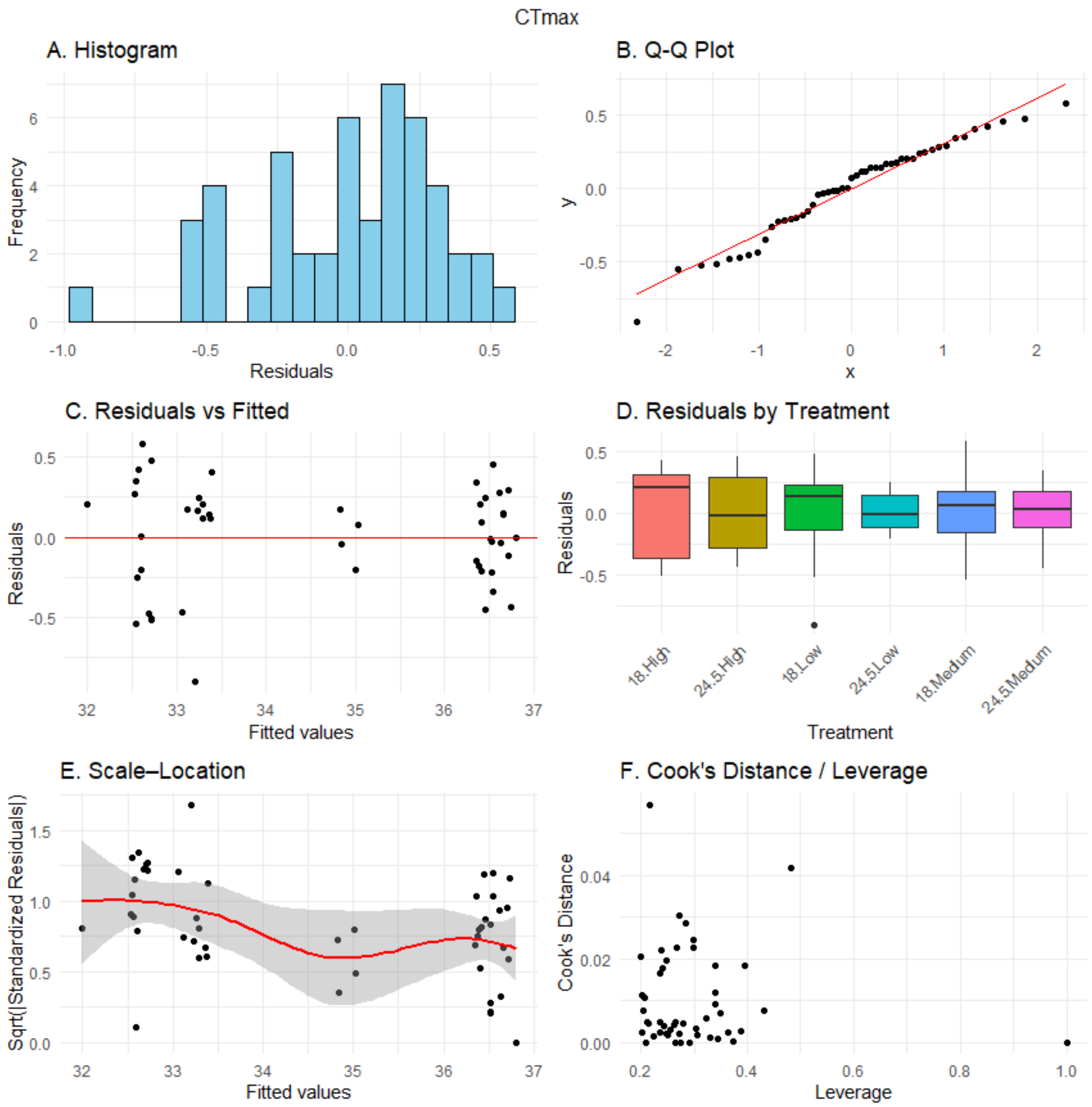


**Figure S3**. Null model assumptions of normality and homoscedasticity of heat tolerance ($CT_{max}$ in °C) were evaluated using standard diagnostic plots: (A) histogram of residuals, (B) Q–Q plot, (C) residuals vs. fitted values, (D) residuals by treatment (Temperature × Food_quality), (E) scale–location plot, and (F) Cook's distance versus leverage plot.

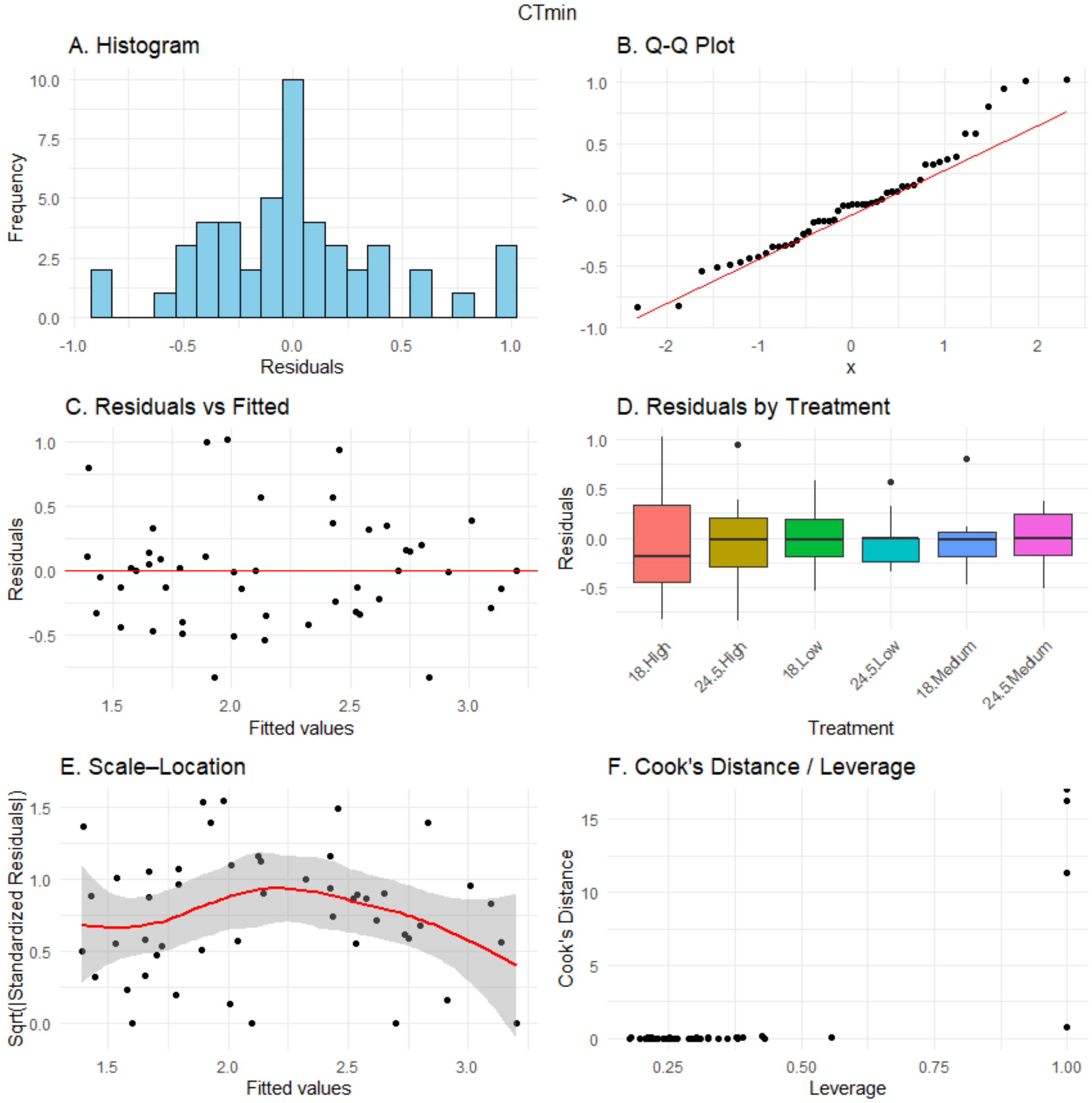


**Figure S4**. Null model assumptions of normality and homoscedasticity of cold tolerance ($CT_{min}$ in °C) were evaluated using standard diagnostic plots: (A) histogram of residuals, (B) Q–Q plot, (C) residuals vs. fitted values, (D) residuals by treatment (Temperature × Food_quality), (E) scale–location plot, and (F) Cook's distance versus leverage plot.

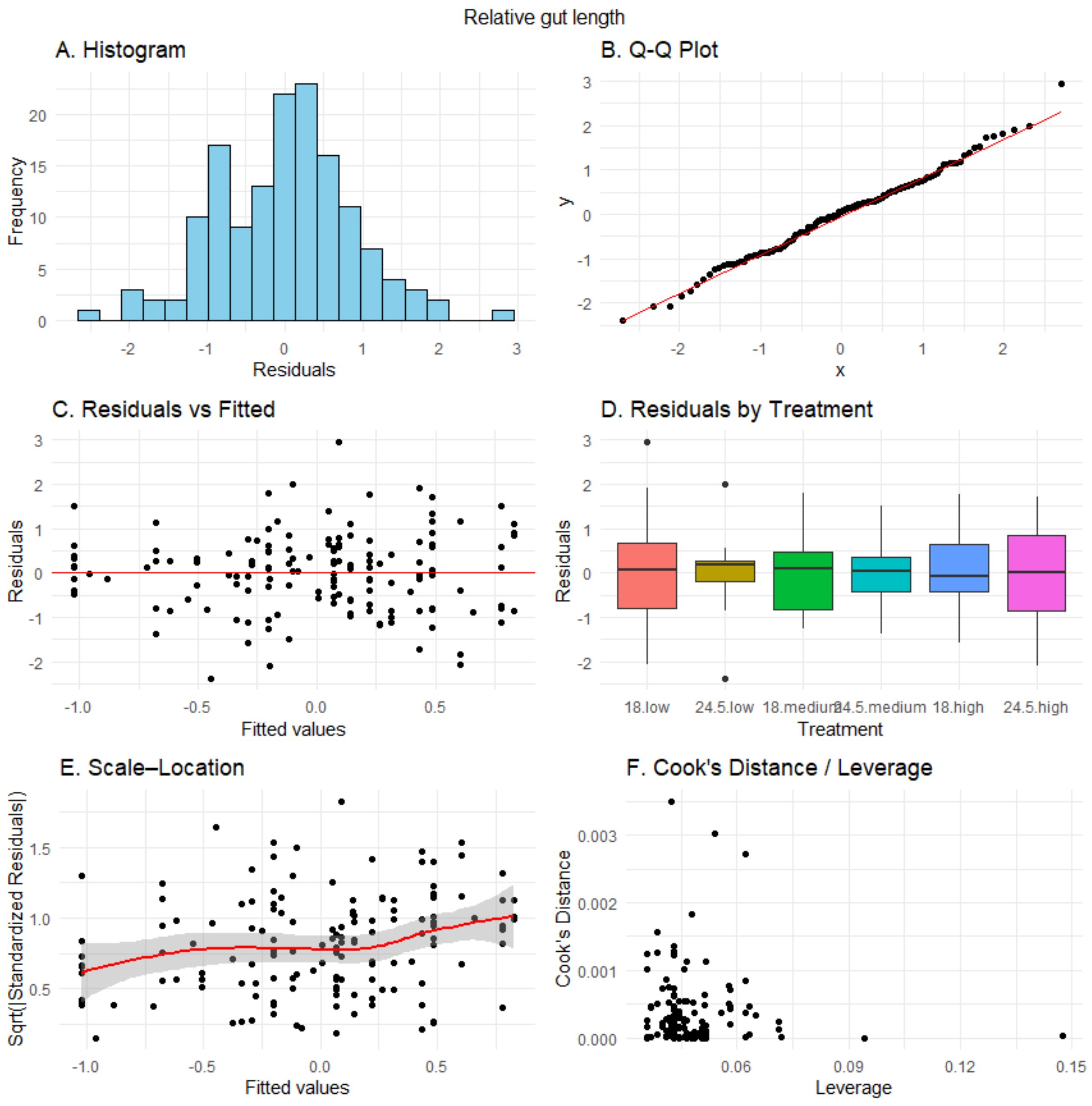


**Figure S5**. Null model assumptions of normality and homoscedasticity of relative gut length were evaluated using standard diagnostic plots: (A) histogram of residuals, (B) Q–Q plot, (C) residuals vs. fitted values, (D) residuals by treatment (Temperature × Food_quality), (E) scale–location plot, and (F) Cook's distance versus leverage plot.

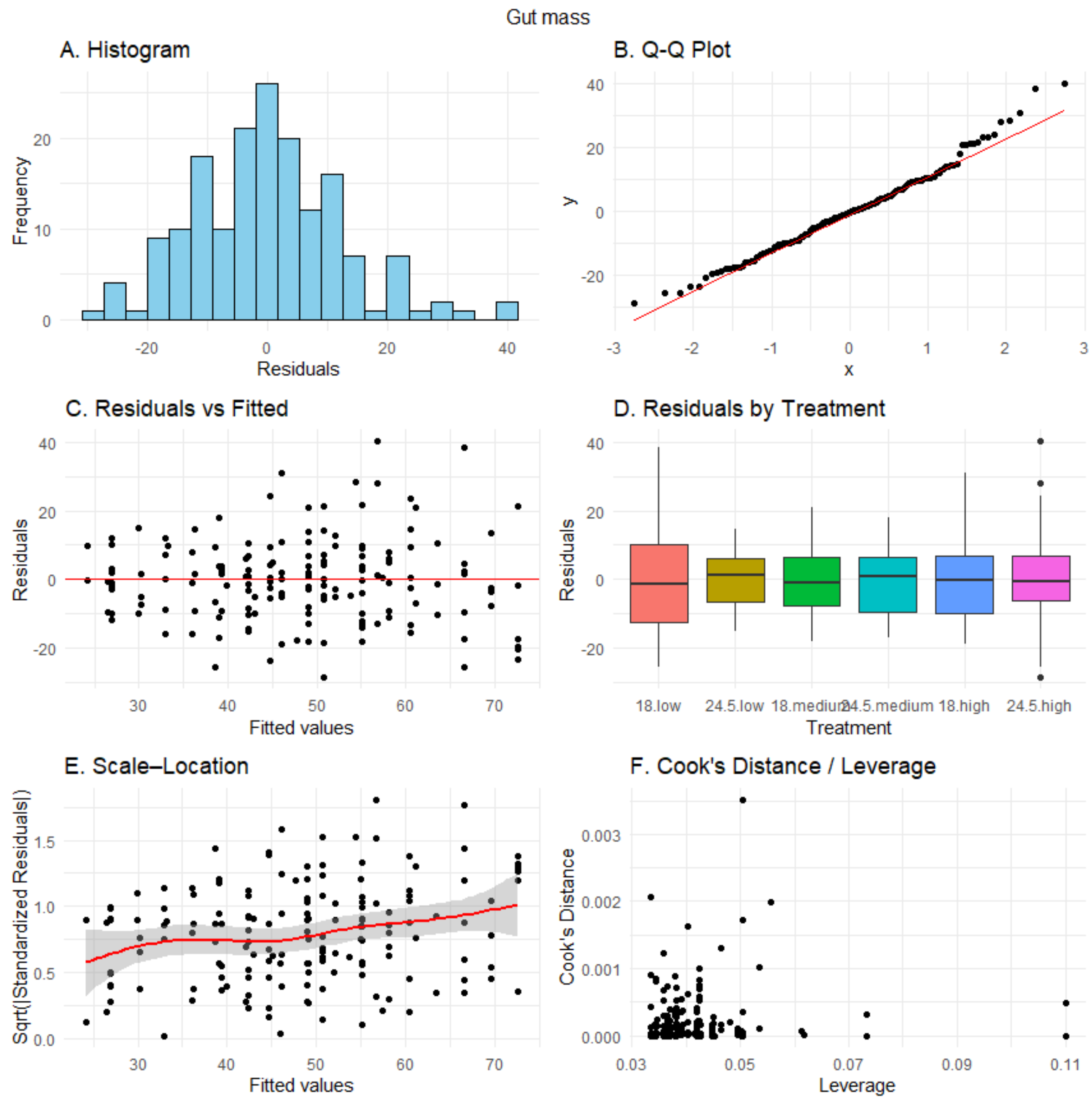


**Figure S6**. Null model assumptions of normality and homoscedasticity of gut mass were evaluated using standard diagnostic plots: (A) histogram of residuals, (B) Q–Q plot, (C) residuals vs. fitted values, (D) residuals by treatment (Temperature × Food_quality), (E) scale–location plot, and (F) Cook's distance versus leverage plot.

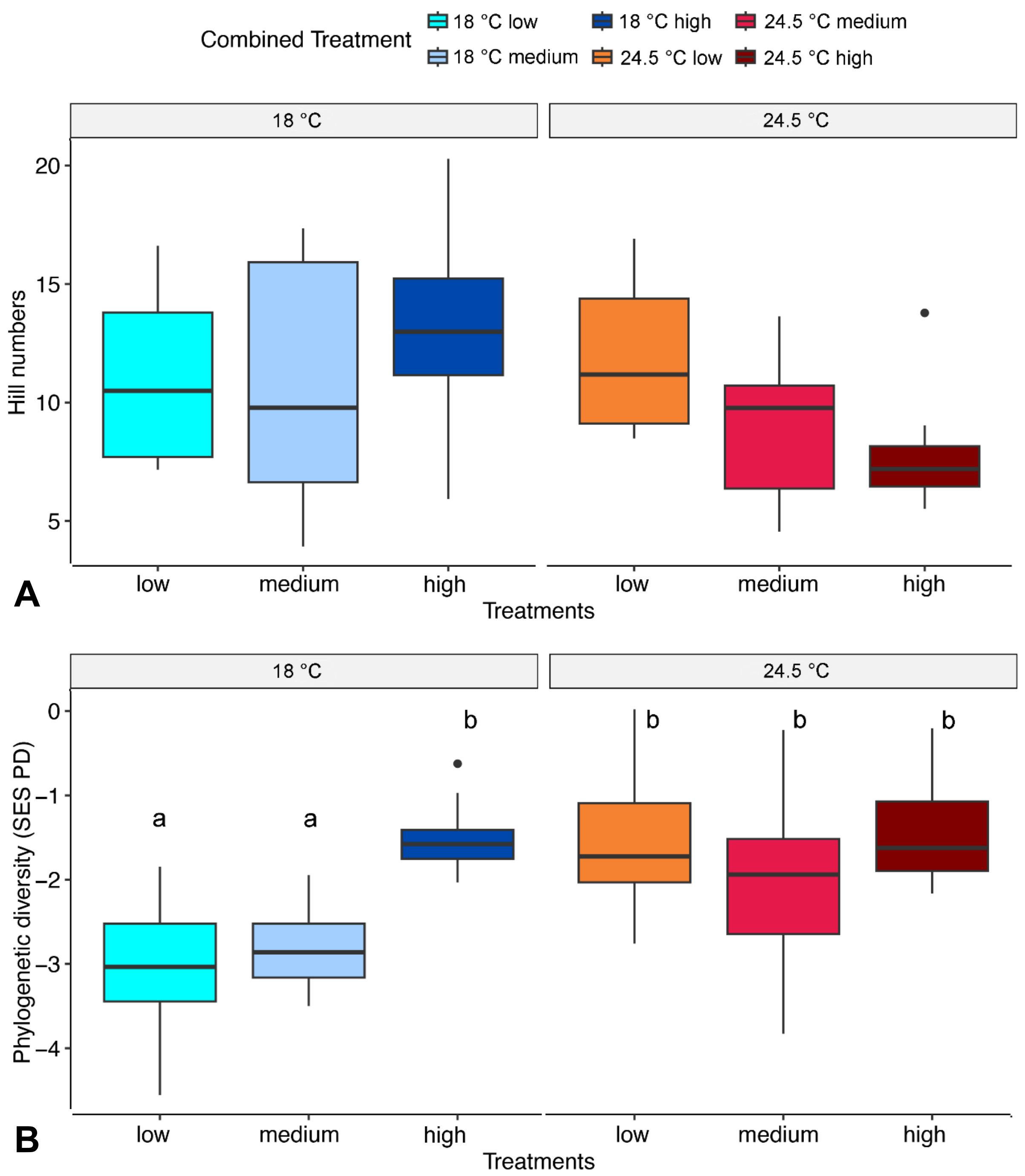


**Fig. S7**. Effects of food quality and rearing temperature on gut bacterial diversity in larvae of *Rana temporaria*: (A) Hill numbers and (B) Faith's phylogenetic diversity. Food quality is represented by increasing levels of protein, fat, and components of animal origin.

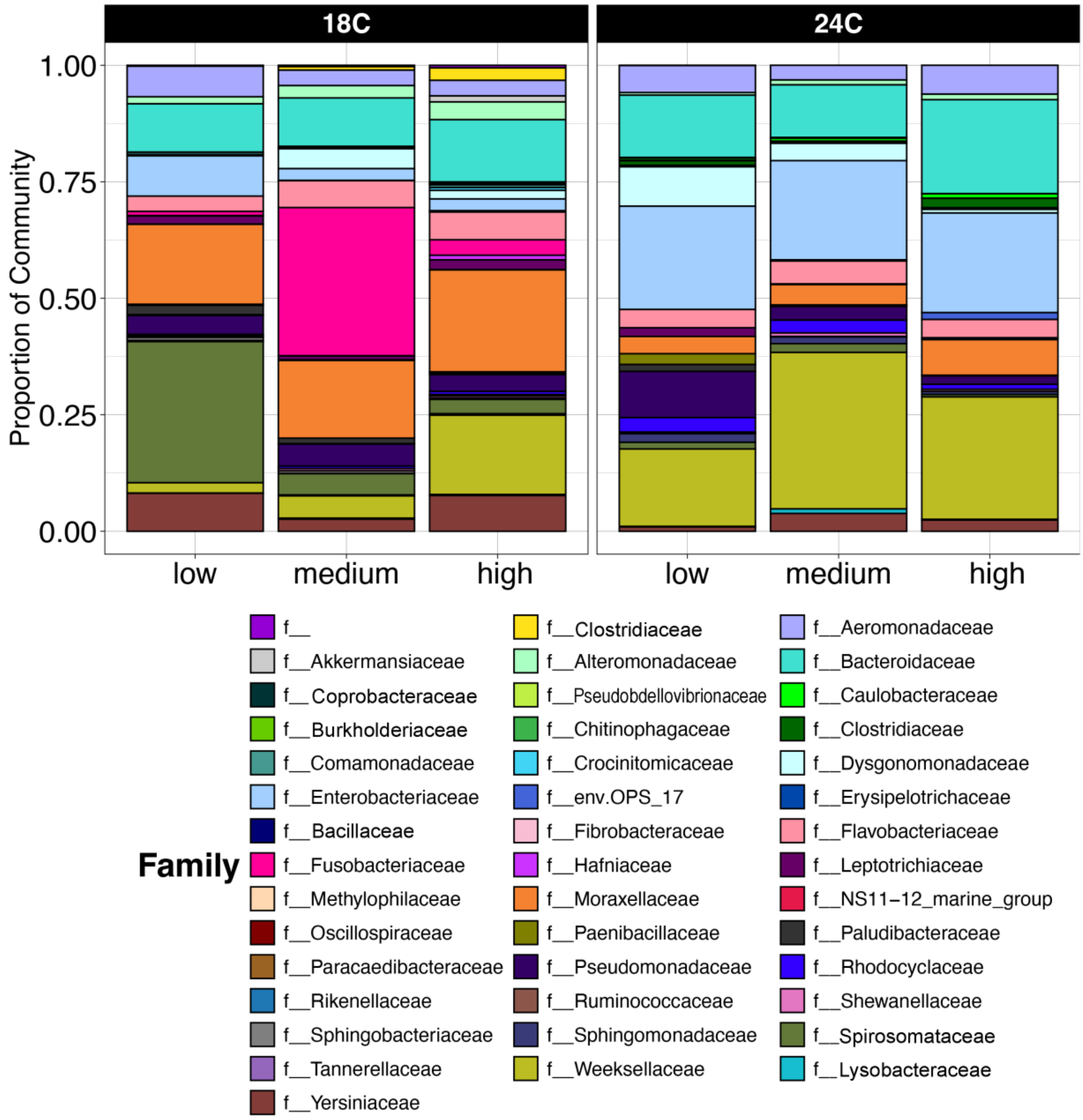


**Fig. S8**. Comparison of microbiome composition at the family level in the guts of larvae of *Rana temporaria* reared under experimental conditions represented by two temperatures (18 °C and 24.5 °C) and three diets with increasing levels of protein, fat, and components of animal origin (considered as low, medium, and high quality food).

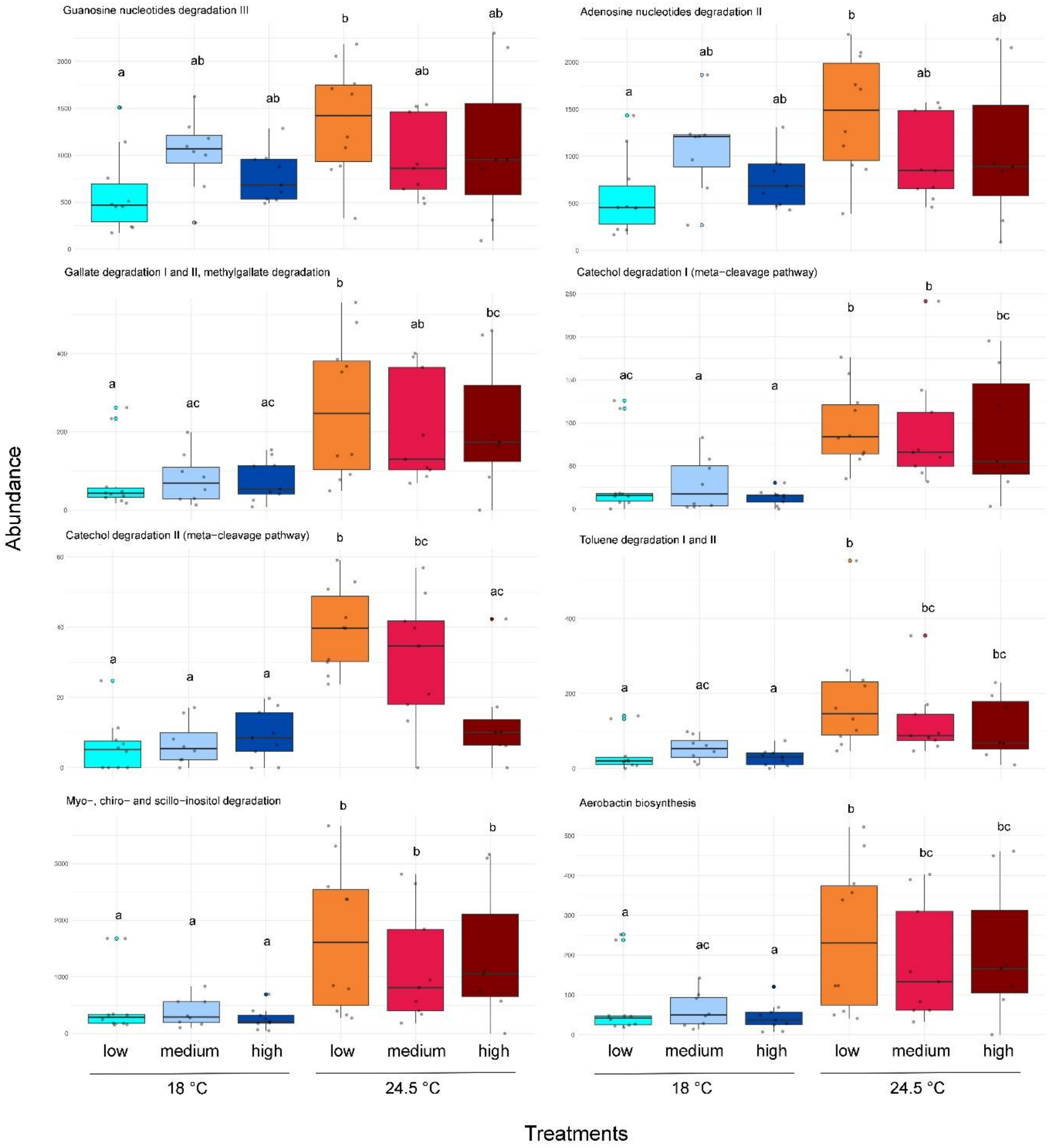


**Fig. S9**. Pathways predicted to increase with temperature in the guts of larvae of *Rana temporaria* exposed to two rearing temperatures and three food treatments with increasing levels of protein, fat, and components of animal origin (considered as low, medium, and high quality food).

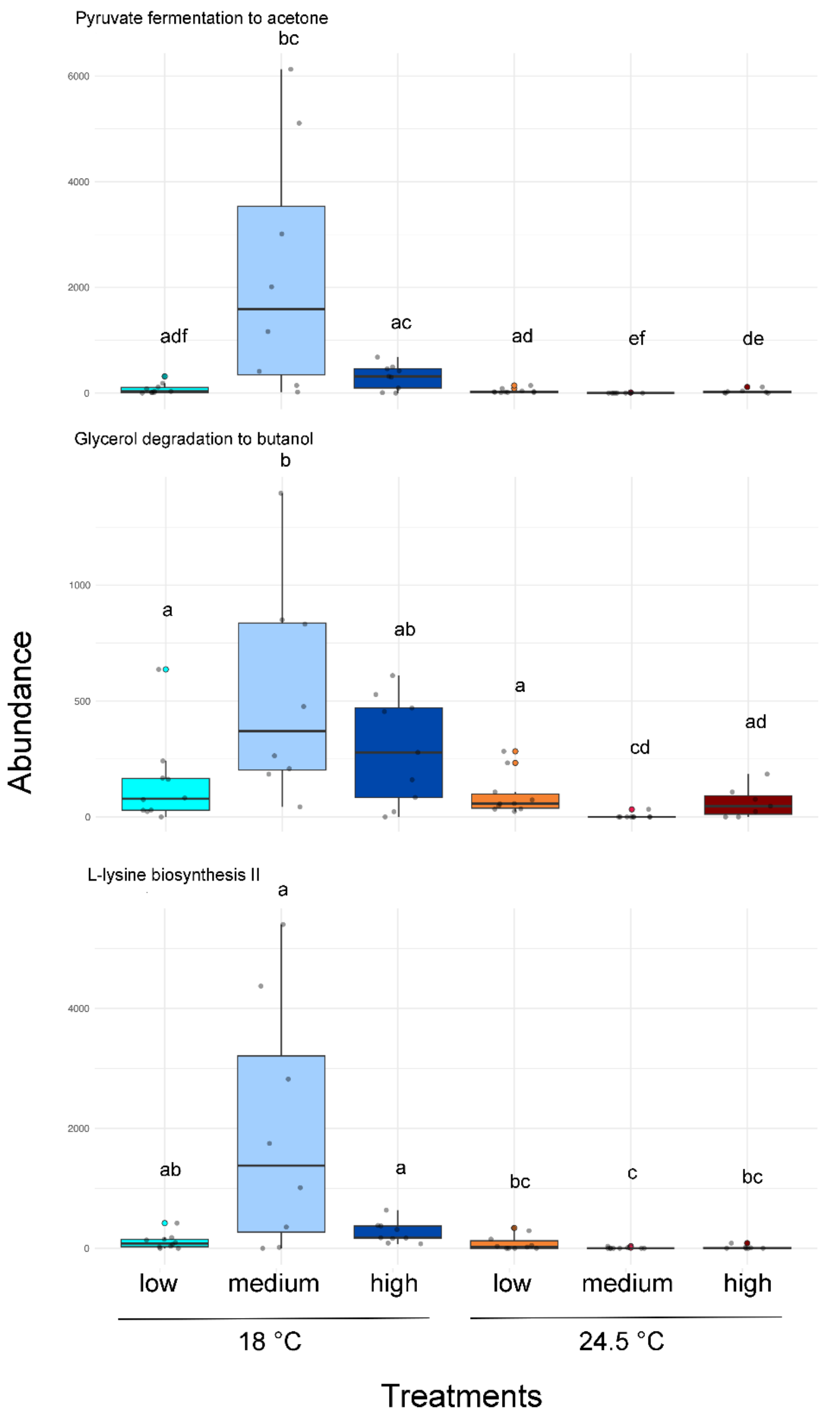


**Fig. S10**. Pathways predicted to decrease with temperature in the guts of larvae of *Rana temporaria* exposed to two rearing temperatures and three food treatments with increasing levels of protein, fat, and components of animal origin (considered as low, medium, and high quality food).
